# Integrated single-cell profiling dissects cell-state-specific enhancer landscapes of human tumor-infiltrating T cells

**DOI:** 10.1101/2022.03.16.484513

**Authors:** Dania Riegel, Elena Romero-Fernández, Malte Simon, Akinbami Raphael Adenugba, Katrin Singer, Roman Mayr, Florian Weber, Charles D. Imbusch, Marina Kreutz, Benedikt Brors, Ines Ugele, Jens M. Werner, Peter J. Siska, Christian Schmidl

## Abstract

Despite extensive studies on the chromatin landscape of exhausted T cells, the transcriptional wiring underlying the heterogeneous functional and dysfunctional states of human tumor-infiltrating lymphocytes (TILs) is incompletely understood. Here, we identify tissue-specific and general gene-regulatory landscapes in the wide breadth of CD8^+^ TIL functional states covering four cancer entities using single-cell chromatin profiling. We map enhancer-promoter interactions in human TILs by integrating single-cell chromatin accessibility with single-cell RNA-seq data from tumor entity-matching samples, and prioritize key elements by super-enhancer analysis. Our results reveal a human core chromatin trajectory to TIL dysfunction and identify involved key enhancers, transcriptional regulators, and deregulated target genes in this process. Finally, we validate enhancer regulation at immunotherapeutically relevant loci by targeting non-coding regulatory elements with potent CRISPR activators and repressors. In summary, our study provides a framework for understanding and manipulating cell-state-specific gene-regulatory cues from human tumor infiltrating lymphocytes.

## Main

Dysfunctional T cells, also termed exhausted T cells (T_EX_), are limited in their ability to clear pathogens and malignant cells. Accordingly, recent research has focused on the mechanisms of T cell dysfunction, which revealed a progressive and partly irreversible reprogramming of the T cells’ chromatin landscape along with impaired effector molecule expression, proliferation, and survival^1^. The chromatin changes are suggested to lock T cells in their dysfunctional state, which represent a hurdle for long-lasting T cell re-invigoration by checkpoint blockade^2^, and impair adoptive cell therapy with chimeric antigen receptor (CAR) T cells^3^. Importantly, several studies identified cells with increased self-renewal capacity in the heterogeneous pool of exhausted T cells, which respond to checkpoint blockade and give rise to terminally differentiated cells that are essential for viral or tumor control^4, 5, 6, 7, 8, 9^. Defining the heterogeneity and regulatory cues in human TILs is therefore crucial to understand and improve responses of immunotherapeutic applications.

On the molecular level, gene-regulatory changes distinguishing dysfunctional from functional T cell states were traced down to exhaustion-specific distal regulatory elements (enhancers) that drive the expression of genes contributing to T cell dysfunction such as the immune-checkpoint Programmed Cell Death 1 (*PDCD1*)^10^. The functional importance of individual enhancer elements in lymphocyte biology is well established, but only known in detail for a small subset of the hundreds of thousands of regulatory regions identified genome-wide. A classic example is the locus control region at the Th_2_ cytokine locus, which orchestrates the cell state-dependent expression of IL-4, IL-5, and IL-13 via long-range chromatin interactions^11^. Signal-dependent functional modulation of T cell states by enhancers was further exemplified by the deletion of one intronic DNA element in the *IL2RA* gene (encoding CD25, part of the high-affinity IL-2 receptor), which shifted the polarization of naïve T cells from induced T_reg_ towards a pro-inflammatory Th_17_ cell state *in vivo*^12^. Another study demonstrated, that the ablation of an enhancer upstream of *PDCD1* (encoding PD1) decreased activation-induced PD1 expression in antigen-specific CD8^+^ T cells, which increased memory T cell formation and improved tumor control *in vivo*^13^. Analogous in the human system, the deletion of distal enhancers of the *PDCD1* gene reduced expression of the co-inhibitory receptor PD1 in CAR T cells during chronic *in vitro* activation^14^, which exemplifies the therapeutic relevance of enhancer editing. Thus, studying cell-state-specific enhancer regulation is critical to understand the molecular mechanisms of immune system processes such as T cell (dys-)function. However, detailed insights into enhancer regulation in human TILs are limited by several obstacles.

First, data on chromatin dynamics upon T cell dysfunction are mainly derived from bulk profiling of murine T cells from cancer or chronic infection models or human healthy donors^1, 2, 10, 15, 16, 17^. Despite computational efforts to unify T cell chromatin data over model systems, differences between chronic infection and cancer models are known^18^. In line with this, recent studies on CAR T cell exhaustion uncovered divergences between murine and human T cell dysfunction-specific regulomes^14^, which is expected due to the reduced conservation of enhancers over species in comparison to protein-coding genes^19^. So far, only limited numbers of studies investigated the single-cell chromatin landscape of human TILs in single cancer entities^20^. Hence, a comprehensive and generalized description of heterogeneous T cell chromatin states and their associated enhancer landscapes in human cancer is missing. Second, there is a striking imbalance between the pure mapping of chromatin changes at distal non-coding elements and our understanding of which target genes are affected. As enhancers can act over long distances, systematic mapping of enhancer-promoter interactions in human tumor infiltrating T cells would be critical to generate hypotheses about the regulation of key genes *in vivo*. Third, as chromatin mapping studies often yield thousands of putative cell-state-specific enhancer elements, prioritization of genes and their connected enhancers is urgently needed to pinpoint central regulatory cues of distinct functional and dysfunctional T cell states that can be targeted to improve cell therapy products including CAR T cells.

In this study we therefore sought to identify the enhancer regulation of key genes in human TILs. We could identify a core gene-regulatory trajectory to T cell dysfunction in tumors by unbiased single-cell analysis of the chromatin landscape of thousands of CD8^+^ TILs derived from several patients over four cancer entities. Integration of entity-matching scRNA-seq data allowed us to predict potential enhancer-promoter links over a wide breadth of functional as well as dysfunctional T cell states, from which we prioritized critical elements by super-enhancer analysis. Finally, we employed CRISPR-based perturbations to validate selected enhancers at top-ranked genes encoding PD1 and TCF1. In summary, our study reveals that, despite tissue-specific remodeling of human TILs, there are general core gene-regulatory patterns underlying T cell functional and dysfunctional states, which can be modulated by enhancer targeting via CRISPR activation/interference. As a perspective, our data provides a framework to understand and manipulate gene expression in human T cells via the non-coding genome in the context of cancer.

## Results

### Single-cell chromatin landscapes over a wide breadth of human CD8^+^ T cell states

To investigate the chromatin landscape of human TILs we first adapted a published plate-based single-cell ATAC-seq assay^21^. We optimized cell lysis buffer conditions of current protocols^21, 22^ to maximize signal-to-noise ratio, and verified a low doublet frequency in single-cell sorting by performing species-mixing experiments of human Jurkat and murine 3T3 cells (**Figure S1A-C**). Aggregated single-cell chromatin profiles from our plate-based assay closely recapitulate profiles of matching bulk ATAC-seq profiles, confirming our experimental conditions (**Figure S1D-E**). Based on this protocol, we isolated T cells and sorted roughly equal numbers of CD8^+^PD1^+^ as well as CD8^+^PD1^-^ TILs per experiment from human head and neck squamous cell carcinoma (HNSCC, n=4 patients, total of 13 samples), hepatocellular carcinoma (HCC, n=4 patients, total of 13 samples), clear cell renal cell carcinoma (ccRCC, n=4 patients, total of 14 samples), and tumor-adjacent control tissue from ccRCC and HCC (six samples from four patients in total) and subjected them to plate-based scATAC-seq (**Figure 1A**, gating and re-analysis **Figure S1F-G**). As additional controls, we performed plate-based scATAC-seq on healthy donor PBMC-derived CD8^+^ T cells with and without *in vitro* activation (n=5 donors, total of 9 samples; all cancer patients and healthy donors see **Table S1**). We further included parts of one publicly available dataset of droplet-based scATAC-seq of whole T cells from patients with basal cell carcinoma undergoing immunotherapy (BCC, n=4 patients, total of 7 samples) as well as sorted naïve and memory PBMC-derived control CD8^+^ T cells from a diabetes study^20, 23^. Quality control of the combined dataset confirmed a high enrichment of open chromatin fragments at transcription start sites (TSS) and unique fragments across the combined (**Figure 1B**) as well as in individual samples (**Figure S2A**). Sequencing libraries revealed the expected ATAC-seq fragment size distributions and enrichment patterns at transcription start sites (TSS), further validating the quality of scATAC-seq of TILs from primary cancer specimen (**Figure S2B-C**). Our goal is to provide a comprehensive transcription-regulatory view over diverse functional and dysfunctional T cell states. Therefore, we examined the sequenced samples for surface markers and cytokine expression capacity by flow cytometry. Representative sample analysis shows that HCC and ccRCC TILs have lower frequencies of cytokine-secreting cells than corresponding T cells from tumor-adjacent tissue controls or activated PBMCs, indicating functional impairment of TILs (**Figure 1C**, gating **Figure S1H**). In contrast, HNSCC TILs had a high cytokine secretion, indicating greater TIL functionality compared to the other investigated entities. Systematic flow cytometry analysis of all samples included in scATAC-seq profiling confirms a functional gradient of T cells ranging from severely dysfunctional (high PD1 surface protein expression in combination with low TNF/IFNγ production) to highly functional (low PD1 MFI with a high percentage of TNF/IFNγ double-positive cells) T cell populations (**Figure 1D**). In control CD8^+^ T cells from PBMCs, PD1 MFI showed a slight positive correlation with cytokine production, while in *ex vivo* TILs this correlation was negative, indicating functional impairment upon chronic activation in the tumor environment. In summary, we generated high-quality single-cell chromatin profiles from human T cells spanning a wide breadth of functional and dysfunctional states.

**Figure 1:**
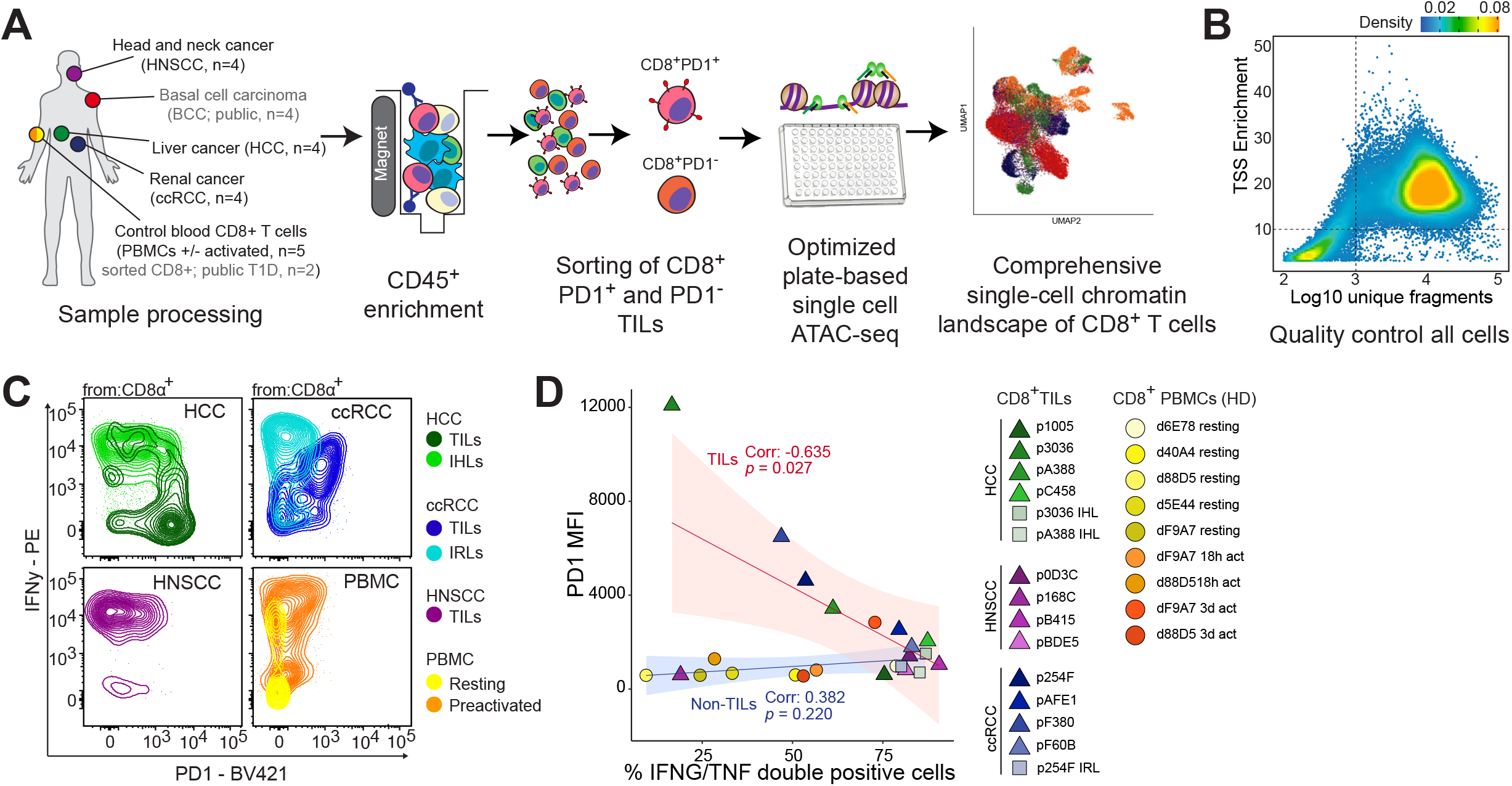
ScATAC-seq workflow and flow cytometric analysis of patient samples. (A) Schematic overview of the experimental setup and sample processing. (B) Transcription start site (TSS) enrichment and number of unique fragments from each single-cell in the ATAC-seq data. Dashed lines indicate cut-offs to discard cells before downstream analysis. (C) Exemplary flow cytometry contour plot showing PD1 vs IFNγ expression of CD8^+^ T cells derived from different tumor entities, tissue controls (IHL= intrahepatic lymphocytes; IRL= intrarenal lymphocytes), and blood. (D) Summary FACS data obtained from samples analyzed by scATAC-seq. PD1-MFI (normalization PD1 negative MFI subtracted from PD1 positive MFI) vs % IFNγ and TNF double-positive cells is shown. Correlation testing was performed using the Pearson coefficient. Gating is shown **in Figure S1H**.

To focus the analysis strictly on CD8^+^ TILs we first removed chromatin profiles of non-CD8^+^ T cells contained in the publicly available BCC sequencing data based on the chromatin accessibility profiles and gene activity scores of the *CD8A, CD8B* and *CD4* genes (**Figure S2D-E**). The integrated, combined data set of all CD8^+^ cells (n=26,479) was then subjected to batch effect correction, dimensionality reduction, clustering, and UMAP embedding for visualization purposes (**Figure 2A**). Based on the genome-wide chromatin accessibility profiles we identified a total of 11 distinct clusters representing different CD8^+^ T cell states that assembled from different entities, patients, *in vitro* or *in vivo* treatment states, cell sorting markers, samples, and scATAC-seq technologies (**Figure 2A-C, Figure S3A-D**). We next aimed to dissect the different cell states that were present in our broad sample collection and identified a total of 8,453 unique marker genes over all clusters, exposing wide-spread chromatin reprogramming among different CD8^+^ T cell subpopulations (**Figure 2D, Table S2**). Gene activity of cluster-specific markers and selected genes (**Figure 2E**) demonstrate that cluster 7 comprised mucosa-associated invariant T cells (MAIT cells, high gene activity of *ZBTB16*). Cluster 6 contains naïve CD8^+^T cells from blood (high gene activity of *LEF1*; in addition, naïve-sorted cells almost exclusively populated this cluster, see **Figure S3B**), while *in vitro* activated PBMC-derived CD8^+^ T cells profiled by scATAC-seq constituted mainly cluster 5 (high activity of *AHRR*; sample origin is almost exclusively from *in vitro* activation experiments, see **Figure S3A**). Cluster 11 displayed high activity of memory T cell marker genes including *IL7R*, while cluster 9 (and partly cluster 8) resembled cytotoxic effector cells (*CX3CR1* activity). Clusters 8 and 3 were characterized by considerable *NR4A1* as well as heat shock protein gene activity (*HSP90AA1*), suggesting recent *in vivo* TCR activity and a heat-shock response. While cluster 10 was enriched for exhausted precursor T cell (T_PEX_) marker genes (*CXCR5, TCF7*), clusters 1-4 all showed a high activity of T cell dysfunction markers including *HAVCR2* (encoding for TIM3), *ENTPD1* (encoding for CD39), and *TOX*. Indeed, when inspecting the genetic locus of *TOX*, a key driver of T cell dysfunction^24, 25, 26^, we observed a vast increase in chromatin accessibility in aggregated single-cell signals in clusters 1-4 (**Figure 2F**). We then projected broader CD8^+^ T cell signatures from public gene-expression datasets to better annotate the T cell clusters (**Figure 2G, Figure S3E**). Signature enrichment analysis confirmed MAIT programs in cluster 7, and naïve T cell signatures in cluster 6. Enrichment of cytotoxic T cell signatures were observed predominantly in cluster 9^27, 28, 29^. In contrast, clusters 1-4 were enriched in genes associated with dysfunctional T cells from melanoma^29^. Here, a similar enrichment was observed for genes from a validated exhaustion signature, which was generated based on chromatin and expression data and validated by MassCytometry^30^, further confirming dysfunctional T cell states in clusters 1-4. These clusters also displayed high activity of genes found in TILs from melanoma patient that did not respond to checkpoint therapy with an anti-PD1 antibody^5^. Of note, HNSCC-derived cells were underrepresented in dysfunctional clusters (**Figure S3F-G**), which is in line with flow cytometry analysis that placed TILs from HNSCC on the functional end of the spectrum regarding cytokine production (**Figure 1C-D**). Contrary to dysfunctional clusters, cluster 11 was enriched with genes found in CD8^+^ T cells of responders to checkpoint blockade^5^. In summary, our data provide unbiased single-cell chromatin accessibility maps over a wide breadth of human primary T cell states that cover immunotherapy-relevant populations, and demonstrate a profound heterogeneity in human TIL chromatin profiles.

**Figure 2:**
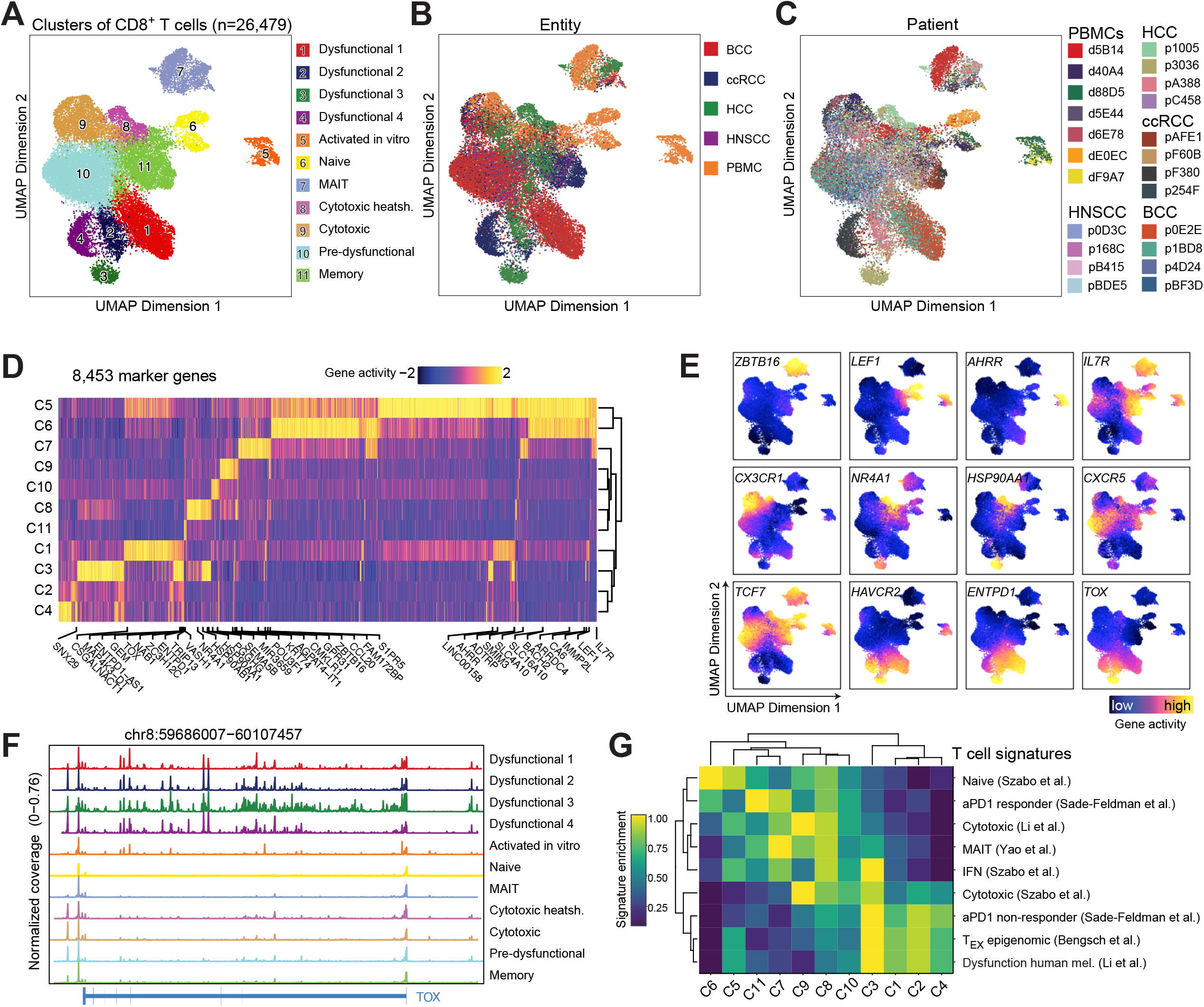
Single-cell ATAC-seq identifies chromatin states over a wide breadth of human primary CD8^+^ T cell functional states. (A-C) UMAP projection of 26,479 single-cell chromatin profiles passing quality control. UMAP projection is colored by clustering (A), entity (B), or patient (C). See also **Figure S3**. (D) Heatmap of gene activity scores determined by ArchR^37^ of all marker genes from each cluster. Data is normalized (Z-Score). Statistical test for marker detection: Wilcoxon, FDR <= 0.01 & log_2_FC >= 0.58. (E) UMAP projection of gene activity of selected genes. (F) Pseudo bulk ATAC-seq profile of each cluster at the *TOX* gene locus. (G) Enrichment of specific T cell signatures in scATAC-seq clusters from **Figure 2A**. Data was normalized using Empirical Percentile Transformation. See also **Figure S3**.

### General and tissue-specific gene-regulatory programs of human TIL states

We next explored gene-regulatory landscapes of the different T cell states. As a basis for analyzing cell-state-specific enhancers and promoters we identified a total of 200,015 unique peaks (accessible chromatin regions) over all 11 clusters, of which 56,894 represented marker peaks for individual clusters (**Figure 3A, Figure S4A, Table S3**). Cluster-specific transcription factor binding motif assessment yielded significant enrichment of TCF motifs in PBMC-derived naïve and *in vitro* activated cells (**Figure S4B**). MAIT cell-specific open chromatin sites were enriched for RORγt binding sites. Besides this, octamer (POU) motifs were enriched in gene-regulatory elements of dysfunctional TIL clusters, however, the contribution of transcription factors from this family to T cell dysfunction is currently unknown. We next evaluated transcription factor motif activity over all >200k open chromatin regions on the single-cell level using chromVAR^31^ and ranked TF motifs accordingly (top 100 motifs, **Figure 3B**). Among the motifs that accounted for the greatest deviation in accessibility over the whole scATAC-seq dataset we identified binding sites of TF associated with T cell subsets, differentiation, activation, and exhaustion, including AP-1, BATF, NFκB, RORγt, T-bet, Nur77 (encoded by *NR4A1*), TCF, and NFAT. Clustering, UMAP visualization of TF deviations, and transcription factor footprinting (**Figure 3C-E, Figure S4C, Table S3**) confirmed RORγt activity in MAIT cells, and TCF activity in naïve T cells. We observed high IFR:BATF motif activity in *in vitro* activated but also to some degree in dysfunctional T cell clusters, while high STAT5 motif activity was characteristic only for activated CD8^+^ T cells from PBMCs. In addition, and in line with reports on regulatory circuits in dysfunctional T cells^32^, we observed prominent activity of NFκB and Nur77 in T_EX_ clusters 1-4. The motif activity of Nur77 was exclusively dominant in dysfunctional T cells, with further motifs of nuclear receptor TFs enriched in these populations (e.g., Esrbb motif). In addition, we observed a loss of KLF as well as ETS:RUNX composite motif activity in dysfunctional T cells, factors that remained unexplored in this context so far. Finally, we discovered a strong heat-shock factor regulatory activity in the HCC-dominated clusters 3 and 8 (HRE.HSF motif activity, **Figure 3C-E**), a notion that was also supported by the presence of HSP90AA1 and HSP90AB1 among the top marker genes in these clusters (**Figure 2D**). Collectively, our data revealed known and potential new regulatory cues in human CD8^+^ T cell states, highlighting a so far unrecognized severe impact of heat shock responses on distinct *in vivo* TIL chromatin states.

**Figure 3:**
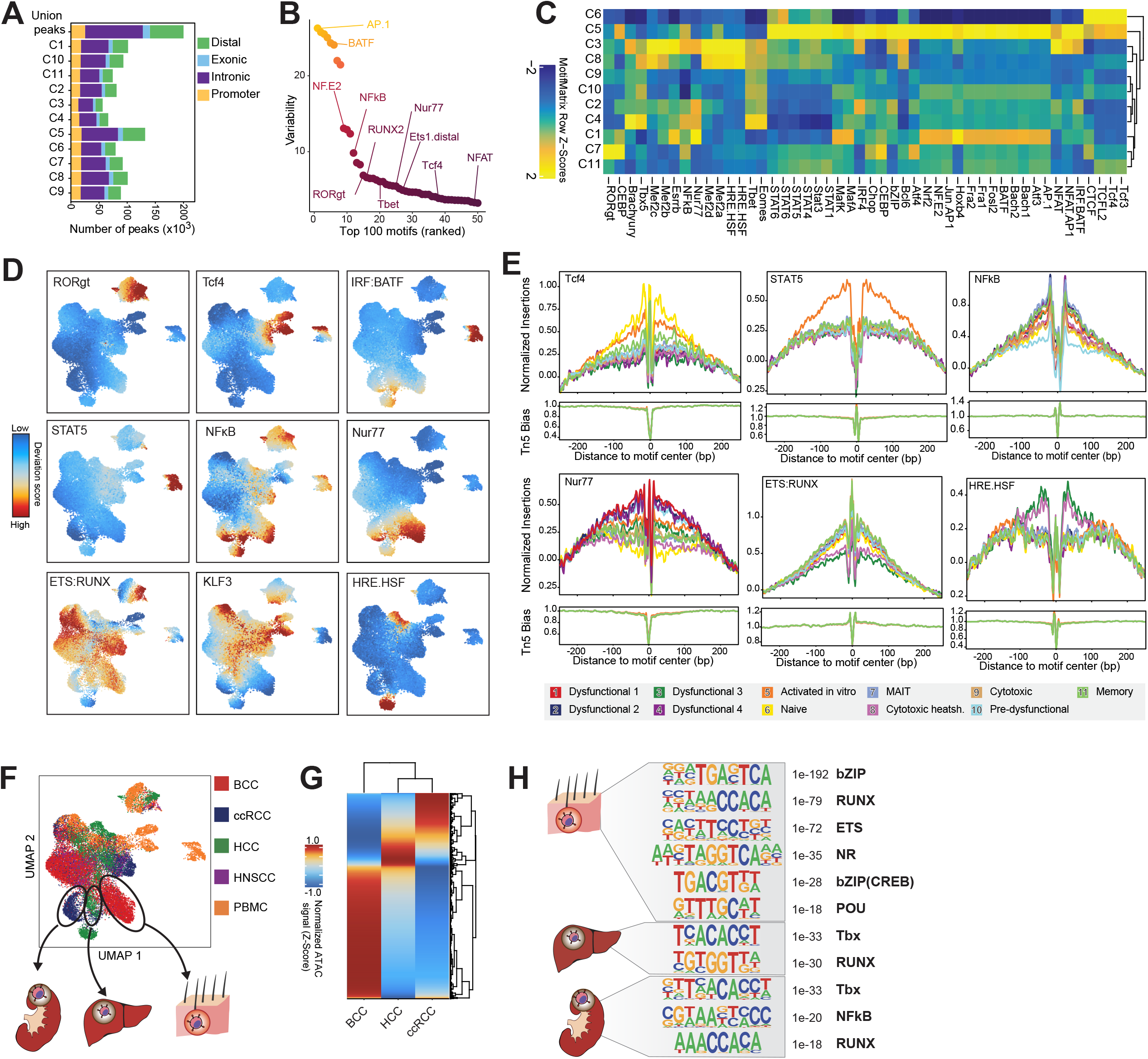
Differential state-specific gene-regulatory cues of human CD8^+^ T cells states. (A) Counts and genomic annotation of accessible chromatin regions detected in the dataset. (B) HOMER transcription factor motif variability detected by chromVAR. Displayed are the top 100 motifs ranked based on their variability over all accessible chromatin regions. (C) Heatmap of motif deviations over the whole ATAC-seq dataset. Shown are motifs with an FDR <= 0.001 (Wilcoxon test) and a mean difference of >= 0.07. (D) UMAP projection of HOMER transcription factor motif deviations on a single-cell level. (E) Transcription factor footprints of selected motifs. Shown is the Tn5 insertion preference-normalized ATAC-seq signal for each cluster at the peaks harboring the indicated TF motif. See also **Figure S4C**. (F) Scheme illustrating analysis of renal, liver, and skin-specific chromatin accessibility profiles of dysfunctional TIL clusters. (G) Accessibility (Z-Score normalized) of tissue-specific peaks in BCC, HCC, and RCC clusters highlighted in Figure (F). (H) *De novo* motif discovery in tissue-specific accessible chromatin regions. Shown is the *de novo* motif, *p*-value (hypergeometric test), and closest matching known motif (or motif family).

We and others have shown that the residing tissue directly shapes the chromatin landscapes of immune cells^33, 34^. Closer inspection of dysfunctional clusters 1, 2, and 4 revealed that they were dominated by cells from a certain cancer tissue type (cluster 1 BCC; cluster 2 HCC; cluster 4 ccRCC, **Figure 3F** and **Figure S3G**), which prompted us to use this information to detect cancer-tissue-specific differences in the chromatin landscape of dysfunctional TILs. We therefore identified marker peaks for each of the three clusters, using the remaining two clusters as background. Additional filtering of peaks that occurred more than once in any of the comparisons yielded 6,332 peaks specific to an entity-dominated cluster (**Figure 3G** and **Table S3**, Cluster 3 was excluded due to the unique heat-shock signature). We then investigated enriched transcription factor binding motifs in cluster-specific peaks. We identified enriched basic Zipper (bZIP), ETS, nuclear receptor (NR) as well as octamer (POU) motifs in the BCC-dominated cluster, T-box (Tbx) and RUNX motifs in both the RCC and HCC clusters, and NFκB motifs in the ccRCC-dominated cluster (**Figure 3H**). Of note, clusters 9, 10, and 11 were more heterogeneously composed of T cells from different entities. Especially cluster 9 (cytotoxic T cells) was composed of similar frequencies of TILs from HCC, BCC, and ccRCC (**Figure S3F**). As these clusters contained regulatory programs associated with functional T cells, our observations indicate an increasing impact of the tissue on TIL chromatin landscapes with progressive T cell dysfunction. In summary, our data suggests tissue/entity-specific chromatin remodelling of TILs with increasing diversification of chromatin patterns in dysfunctional T cells.

### A human core CD8^+^ trajectory to T cell dysfunction in tumors

We next sought to utilize our diverse dataset to identify a unifying pathway to terminal TIL dysfunction in the human system. To this end, we removed PBMC samples and MAIT cells to focus on conventional CD8^+^ T cells isolated from solid cancer specimens and adjacent control tissue, retaining 47 samples from 16 patients. We corrected tissue effects on TILs by differential scaling and by harmonizing data for tissue origin. Clustering identified a total of 6 clusters with 2488 marker genes across all populations (**Figure 4A, Table S4)**. Next, we assessed multiple sources of information to assign functional T cell states to these clusters and identify a chromatin accessibility-based core CD8^+^ trajectory to dysfunction. First, due to cell sorting of PD1^+^ and PD1^-^ populations our data contains the single-cell information on PD1 surface protein expression (**Figure 4B**), which showed that cluster 1 and 2 consisted completely of PD1^+^ cells indicating activation. In contrast, cluster 6 contained mostly PD1^-^ cells, while clusters 3-5 were mixtures of PD1^+^ and PD1^-^ cells. Second, we assessed the distribution of lymphocytes that we profiled from adjacent non-tumorous control tissues (6 samples in total from liver and renal tissue) in the UMAP projections (**Figure 4C**). T cells from control tissue were almost absent from clusters 1 and underrepresented in cluster 2, implying exclusiveness of cluster 1 for tumor-infiltrating T cells. Third, we investigated the activity of established T cell state marker genes (**Figure 4D**) and observed the highest activity of T cell dysfunction markers in clusters 1 and 2 including *PDCD1, HAVCR2, TOX, LAYN, CTLA4*, and *ENTPD1*. Precursor exhausted T_PEX_ marker genes including *TCF7* (encoding TCF1) and *CXCR5* were most active in cluster 3, while memory markers including *IL7R, BACH2* and *LEF1* were observed in cluster 4. We identified cytotoxicity-associated genes in cluster 6 including *PRF1, GZMB, FGFBP2*, and *CX3CR1*, whereas heat shock response genes (*HSP90AA1, HSP90AB1, DNAJB1*) were prevalent in cluster 5. Cluster 5 was also marked by high *NR4A1* gene activity suggesting immediate TCR signalling. These observations imply that cluster 1 and 2 contain dysfunctional cells over all entities with cluster 1 representing the probably strongest TIL dysfunction, and cluster 2 containing proliferating cells (*MKI67* activity). Fourth, we projected curated public gene signatures of T cell states^5, 8, 18, 27, 29, 30, 35^ on the unified core TIL clusters (**Figure 4E**). Specifically, cluster 1 shows signatures of T cell exhaustion identified in human and murine systems of chronic infection and cancer. In line with the high activity of *MKI67*, cluster 2 was enriched in an exhausted/proliferation signature that distinguished it from cluster 1. Cluster 3 was enriched in previously defined signatures of precursor exhausted cells from models of chronic LCMV infection and B16 melanoma, supporting T_PEX_ potential. In contrast to this observation, a different recently described early exhausted precursor signature from a murine infection model^18^ was not enriched in our T_PEX_ cluster. Cluster 6 uniformly displays a cytotoxicity signature, while cluster 5 (heat shock and activation) was enriched in an IFN response signature. Hence, gene signatures also support a developmental trajectory of progenitor cells to terminally dysfunctional cells from cluster 3 to 1. Fifth, we investigated the shift in TIL states post immune checkpoint blockade, where we observed a proportional increase of cells in cluster 1 after treatment (**Figure 4F**). As exhausted cells were described to increase in numbers upon checkpoint blockade^4, 7, 36^, these observations additionally support a general core T_EX_ trajectory from cluster 3 to cluster 1.

**Figure 4:**
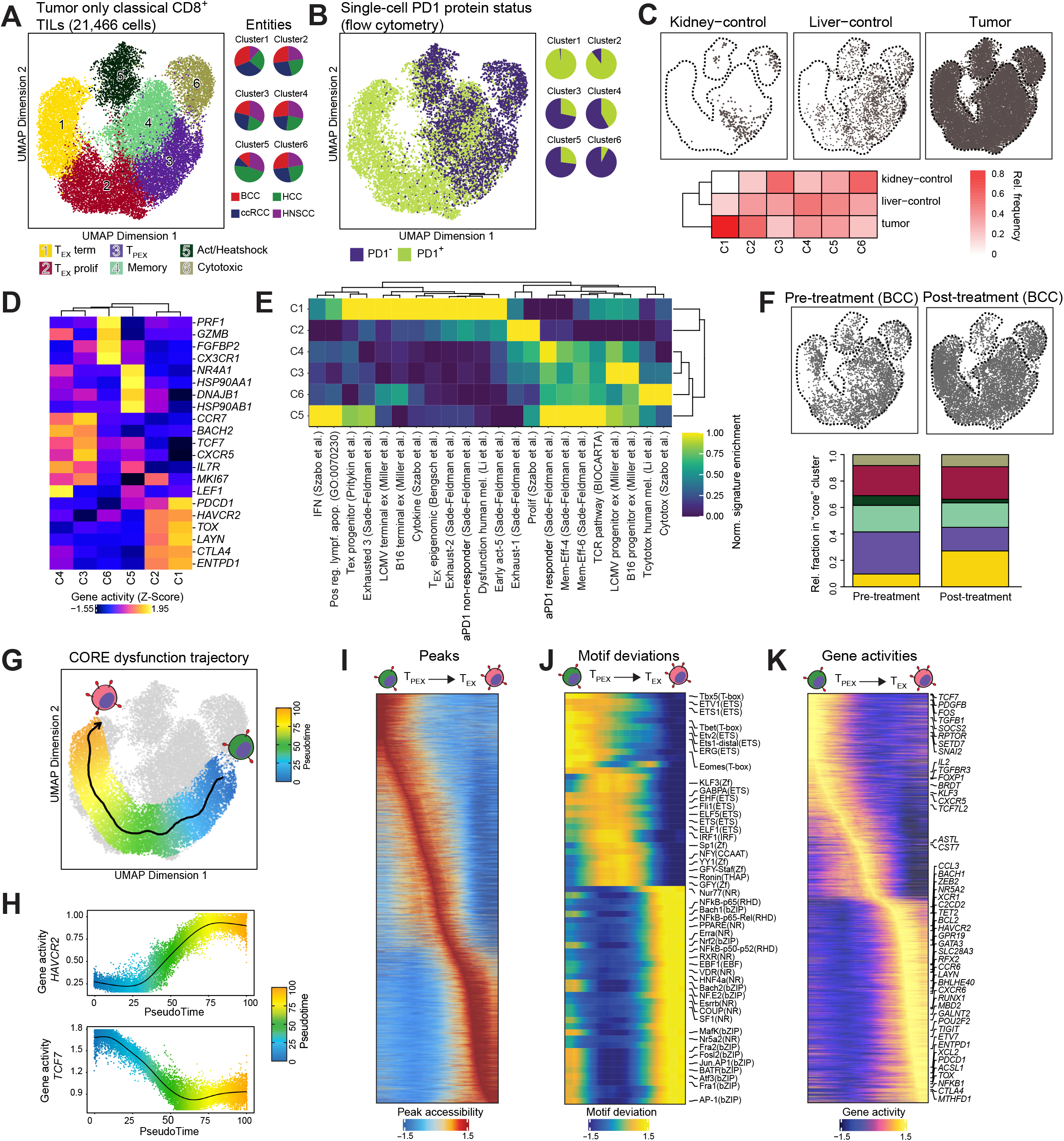
A unifying core pathway to terminal dysfunction of human CD8^+^ TILs. (A) UMAP projection of clustering of 21,466 single-cell chromatin profiles of conventional CD8^+^ T cells from tissues. Relative contribution to clusters (normalized to all cells from the respective entity) is shown in the pie charts. (B) Single-cell PD1 protein expression status (UMAP projection) and relative contribution of PD1^+^ and PD1^-^ cells to each cluster (pie charts). (C) Origin of cells from either adjacent non-tumor kidney control tissue, adjacent non-tumor liver control tissue, or tumor tissue (all entities) in the core TIL UMAP projection (top panel) or represented as a heatmap (cell frequencies normalized to total cells from each origin in the whole dataset, lower panel). (D) Heatmap of Z-Score normalized gene activity scores of selected genes of interest over the TIL core clusters. (E) Enrichment of specific T cell signatures in core TIL scATAC-seq clusters. Data was normalized using Empirical Percentile Transformation (F) Distribution of cells taken from biopsies pre or post immunotherapy (checkpoint blockade) in BCC samples. (G) Core chromatin accessibility pseudotime pathway to CD8^+^ TIL dysfunction. (H) Gene activity of *HAVCR2* (encoding TIM3) and *TCF7* along the pseudotime dysfunction pathway (I-K) Heatmaps of significant dynamic peaks (I), motif deviations (J), and gene activity scores (K) along the T cell dysfunction trajectory.

Indeed, when computing this T_PEX_ to T_EX_ trajectory accordingly (**Figure 4G**), a steady increase of chromatin activity of *HAVCR2* (encoding TIM3) can be observed, while *TCF7* (encoding TCF1) activity is steadily decreasing towards the state of terminal T cell dysfunction (**Figure 4H**). Using this human core exhaustion trajectory, we are able to provide a systematic view of dynamic changes regarding chromatin accessibility, transcription factor activity, and gene activity in human TILs (**Figure 4I-K, Table S4**). Along this trajectory, we could identify several new potential transcriptional regulators and genes that demarked the developmental stages towards T_EX_. As an example, our data suggests differential ETS factor activity between T_PEX_ and T_EX_ (**Figure 4J**). Further, Zinc finger factor binding motifs including KLF are enriched in T_PEX_ but lose the activity in severely dysfunctional TILs that were dominated by binding motifs of bZIP, nuclear receptors, and NFκB transcription factors. Of note, we discovered high activity of nuclear receptors (E.g., VDR, Erra, RXR, Esrrb) as well as EBF among the T_EX_-associated TF binding motifs. Focusing on the pathway to terminal dysfunction, and in agreement with decreased KLF motif activity, we note a reduction of *KLF3* gene activity (**Figure 4K**). In the transition from T_PEX_ to T_EX_, we identified activity loss in several genes including *FOXP1, BRDT, SOCS2, SETD7, SNAI2*, and *RPTOR*. In addition to well-known genes such as *TOX, ENTPD1, LAYN*, and *HAVCR2*, the core signature of severely dysfunctional T_EX_ displayed strong chromatin remodelling and increased gene activity at loci largely undescribed in this context including transcription factors (*ETV7, POU2F2, RFX2, BACH1, ASCL1*), metabolic enzymes and transporters (*MTHFD1, SLC28A3*), epigenetic modifiers (*TET2, MBD2*), intra/intercellular signalling molecules (*GPRR19, CCR6, XCL2, XCR1, CXCR6*), and others. Overall, our unbiased single-cell chromatin accessibility analysis could identify *in vivo* core chromatin landscapes and differentiation pathways of functional and dysfunctional human tumor-infiltrating T cell states.

### Integrated analysis of single-cell chromatin and transcriptional landscapes identifies putative enhancer regulation of TIL genes

Despite comprehensive mapping of gene-regulatory elements in TILs, the lack of information about their interaction with target genes prevents understanding and detailed investigation of gene-regulatory cues. We therefore sought to exploit the possibility to predict enhancer-gene interactions by integrated scRNA-seq and scATAC-seq^37^ to provide a comprehensive resource of human TIL gene regulation. To this end, we processed entity-matching publicly available scRNA-seq data from BCC^36^, HCC^38^, HNSCC^39^, RCC^40^, and resting as well as *in vitro* activated PBMCs^27^ (**Figure S5A**). We removed all but conventional CD8^+^ cells (either by provided annotation or our own classification, see **Methods** for details and **Figure S5B-C**), and clustered the remaining 38,356 CD8^+^ human T cells based on their transcriptomes (**Figure 5A, Figure S6A**). Low-quality cells (cluster 8) that show low abundance of unique mRNA molecules were excluded from further analyses (**Figure S6B**). Transcription of marker genes (**Figure 5B, Table S5**) and genes of interest (**Figure 5C**) suggest a cytotoxic T cell module in cells from cluster 1 (e.g., *CX3CR1, FGFBP2, FCGR3A, KLF2*/*3, TBX21*). Clusters 2 and 5 contained cells with high expression of naïve and memory markers (*IL7R, CCR7, TCF7*), and cluster 6 represented cycling cells (*MKI67*). Surprisingly, in cluster 7 we found genes involved in antiviral responses and strong IFN signalling (e.g., *ISG15, MX1, IFIT3*), which was different from cluster 4 that resembled more the previously defined heat shock / IFN response cluster identified by ATAC-seq. Specifically, cells in cluster 4 displayed comparatively the highest *NR4A1* levels and heat shock transcripts (e.g., *HSP90AA1, DNAJA1*). Cluster 3 expressed dysfunction markers (e.g., *TOX, ENTPD1, HAVCR2, PDCD1, LAYN*), and the adjacent cluster 0 T_PEX_ marker genes (highest CXCR5 expression, and expression of *TCF7, CD27, GZMK*). Hence, similar to the ATAC-seq data, re-analysis of matching scRNA-seq identified diverse but discrete CD8^+^ T cell states from human primary cancer samples. We then integrated the scRNA-seq and scATAC-seq datasets^41^, which directly aligns cells from both datasets by comparing scATAC-seq gene activity with scRNA-seq gene expression (**Figure 5D**). Cells from scRNA-seq clusters that were most abundantly aligned with each of the 11 scATAC-seq clusters showed clear distribution patterns (**Figure 5E**). Specifically, cytotoxic T cells (ATAC cluster 9) were matched with transcriptomes from cluster 1. Naïve, *in vitro* activated, and memory CD8^+^ T cell populations (ATAC clusters 6, 5, and 11, respectively) were highly enriched with alignments to scRNA-seq clusters 5, 6 and 2. As suggested by our chromatin profiles, the heat shock response signature (scRNA-seq cluster 4) was present both in dysfunctional TILs and potential activated cytotoxic cells (ATAC clusters 3 and 8, respectively). Cells from the scRNA-seq T_EX_ cluster 3 closely aligned with the chromatin-defined dysfunctional TILs (clusters 1-4), while potential T_PEX_ cells (ATAC cluster 10) were correlated with transcriptomes from cluster 0. This analysis demonstrates that we can identify matching transcriptional modules for each conventional CD8^+^ T cell cluster identified by scATAC-seq. We next calculated correlations between peak accessibility and gene expression from the integrated dataset, which allows to predict specific enhancer-to-promoter links. In total, we identified 49,902 peak-to-gene links (PGLs, **Figure 5F, Table S5**). Visual inspection of representative marker gene loci illustrates that peak-to-gene links not simply connect enhancers to the nearest gene but predict long-range interactions skipping genes and open chromatin regions in-between (**Figure 5G**). The integrated analysis is easier to interpret than co-accessibility patterns (**Figure S6C**), and is therefore ideally suited to generate hypothesis of enhancer regulation of critical T cell genes. We next asked if our analysis can recapitulate published enhancer-promoter interactions, and therefore compared the predicted interactions at the *IL2RA* locus with experientially validated positive gene-regulatory elements^12^. To this respect, our predictions linked all of the six previously described enhancers correctly to the *IL2RA* promoter (**Figure 5G**). To further investigate if predicted enhancer-promoter-links potentially represent physical chromatin interactions, we calculated the enrichment of peak-to-gene-links in published chromatin conformation capture sequencing (Hi-C) data from different PBMC-obtained immune cells^42^. As expected, peak-to-gene links were more enriched in CD8^+^ T cell Hi-C contacts than in random enhancer-promoter links, and also more enriched in CD8^+^ cell Hi-C contacts compared to Hi-C chromosomal contacts from more distantly related cell types (**Figure 5H**). Taken together, our integrated analysis of matching single-cell chromatin and transcriptome profiles provides a rich resource of putative enhancer-gene interactions in primary human T cell states.

**Figure 5:**
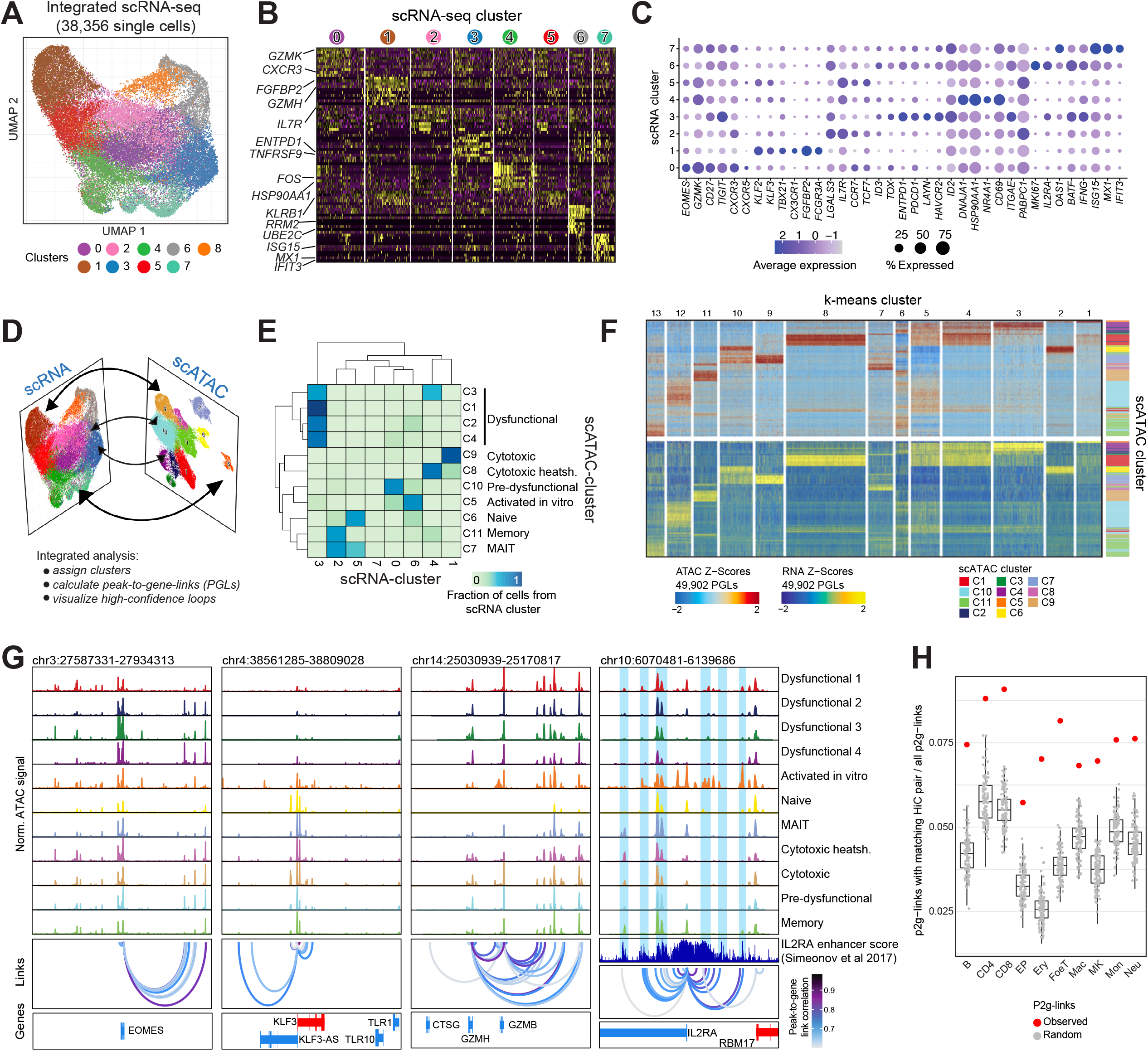
Integrated single-cell RNA and chromatin accessibility profiling predict enhancer-gene interactions of human tumor infiltrating T cells. (A) UMAP projection and clustering of integrated publicly available scRNA-seq covering matching entities used also in scATAC-seq (BCC, RCC, HCC, HNSCC, and CD8^+^ from PBMC with and without activation via anti-CD3/anti-CD28). Related to **Figure S5A**. (B) Heatmap of 10 top markers (selected by highest average log_2_ fold expression change) for each cluster. (C) Dotplot of RNA-expression of selected marker genes and genes of interest from scRNA-seq. (D) Scheme illustrating integration of scATAC-seq with scRNA-seq using the ArchR package. (E) Heatmap of cluster assignment of cells from scRNA-seq to clusters defined in scATAC-seq. Assignments are made based on similarities of gene expression (scRNA-seq) and gene activity score (scATAC-seq). (F) K-means clustered heatmap of peak-to-gene links derived from integration of scATAC-seq and scRNA-seq data. Shown is the accessibility and gene expression of the linked gene, as well as the ATAC-seq clusters. (G) Representative browser plot visualization of pseudo-bulk ATAC-seq signals of all clusters from **Figure 2A**, peak-to-gene-links, and genes at the *EOMES, KLF3, GZMB*, and *IL2RA* loci. Enhancer activity for *IL2RA* mapped by CRISPR-activation^12^ is also displayed. The 6 genomic regions that were classified as *IL2RA* enhancers from that study are highlighted in blue. See also **Figure S6C**. (H) Enrichment of peak-2-gene links in experimentally-derived physical chromatin interaction data from different cell types from publicly available promoter capture Hi-C experiments^42^.

### Perturbation of enhancers using CRISPRi/a validates T cell state-specific gene-regulatory elements at prioritized genes

With the apparent heterogeneity of TILs, a major goal of our study is to delineate functional from dysfunctional TILs to better understand the involved molecular pathways, but also to provide the prospect to manipulate key regulatory elements for immunotherapeutic applications in the future. Therefore, we aimed to prioritize genes and associated enhancers for the identified core cell states. Stretch enhancers, also termed super-enhancers, constitute the accumulation of enhancers that carry an exceptionally high proportion of the active chromatin mark Histone H3 Lysine 27 acetylation, and are enriched at genes that determine cell identity and regulate cell-specific processes^43, 44^. Recently, a method was described to infer super-enhancers centered on chromatin accessibility^45^. The approach is based on counting ATAC-seq peak associations per gene, then ranking the genes based on the number of such specific interactions. We employed this method, but did not use generalized peak-to-gene assignments and instead used our specific peak-to-gene predictions to allocate cluster-specific peaks to their target gene, and then ranked the genes accordingly (**Figure 6A, Table S6**). Using this method of prioritization, we discovered many known key regulators of T cell states (**Figure 6B**). Specifically, among the top-ranked genes in the terminal T_EX_ cluster were *TOX, ENTPD1, HAVCR2, CTLA4, LAYN*, and *BATF*. In the T_PEX_ cluster, our analysis yielded *TCF7* and *CXCR5* among the genes with the most cell-state-specific connected enhancers. Further, we could identify the well-described makers *TBX21, CX3CR1, ZEB2, KLF2*, and *FGFBP2* among the top-ranked genes in the cytotoxic core cluster. Hence, our analysis identifies and prioritizes key mediators of T cell (dys-)functional states. To validate that our resource can identify enhancer-promoter interactions at these key genes we utilized CRISPR-based perturbation approaches. CRISPR activation and interference (CRISPRa and CRISPRi, respectively) can be used to activate or repress distal gene-regulatory elements to evaluate their impact on transcription of target genes^12, 46^. CRISPRa is also able to discover enhancers that are not currently active or inaccessible on the chromatin level, making it a suitable tool to verify functional enhancers in a different cellular context than where the initial enhancer was discovered^47^. We therefore cloned the highly effective ZIM3 CRISPR repressor^48^ and the size-optimized VPRmini CRISPR activator^49^ into a lentiviral vector for stable integration (**Figure 6C**). As a first target, we investigated a putative enhancer for PD1 that was described to be active in exhausted CAR T cells^14^. As PD1 was only expressed at low frequencies in our cells, we tried to increase PD1 expression by enhancer activation with the CRISPRa construct (**Figure 6D**). Indeed, analysis by flow cytometry confirmed a significant increase of PD1-postive cells as well as a trend in increased MFI when targeting the -5 kb enhancer in comparison to a non-targeting (scr) control gRNA (**Figure 6E-F**). We next sought to use the CRISPR tools to validate enhancer regulation at a cell-state-specific gene that distinguishes functional from dysfunctional T cells in the tumor. We therefore browsed peak-to-gene-links as well as core cluster-specific open chromatin regions for genes that were top-ranked in the super enhancer analysis and picked *TCF7* (encoding TCF1) for validation due to its role in exhausted precursors, T cell-memory, and T cell-stemness. As TCF1 was expressed in our model system, we used CRISPRi system for interference with enhancer activity, and targeted a distal putative enhancer that was marked by a cell-state-specific open chromatin region in our analysis. Indeed, although being 27kb distant from the promoter, CRISPRi targeting of this element significantly reduced the frequency of TCF1 positive cells as well as TCF1 protein expression in comparison to a nontargeting control (**Figure 6G-I**). Strikingly, this enhancer was classified as a memory T cell (T_MEM_)-specific enhancer in TILs, which distinguished it from dysfunctional subsets, and shows the usability of our dataset to uncover regulatory elements that may shape T cell functional states in the tumor. In summary, our analysis prioritizes key genes and connected enhancers by integrated single-cell profiling, and we validate regulatory interactions of enhancers and promoters of state-specific TIL genes with CRISPRi/a.

**Figure 6:**
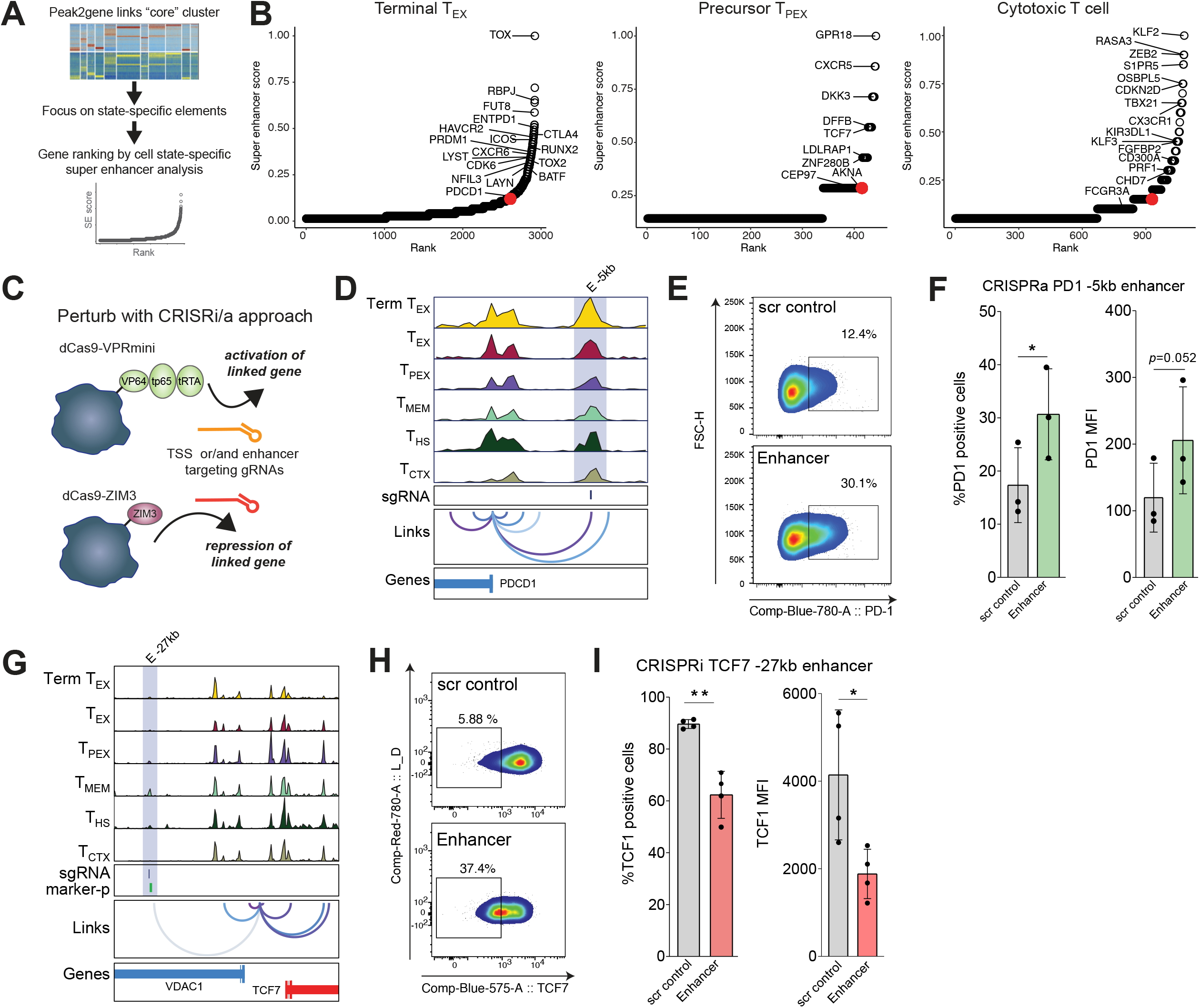
CRISPRi/a validation of cell-state-specific enhancers. (A) Workflow for identification of state-specific enhancers and ranking of connected genes in human TIL cluster. (B) Super-enhancer analysis and ranking of key genes in selected TIL core clusters. (C) Scheme of CRISPR interference and activation approach. (D) *PDCD1* gene locus with pseudo bulk ATAC-seq signal from core clusters with peak-to-gene links. Highlighted is the known enhancer 5 kb upstream of the TSS that is targeted by CRISPRa. (E) Representative flow cytometry plot showing PD1 expression in Jurkat T cells transduced with dCas9-VPRmini and electroporated with a control gRNA or a gRNA targeting the -5kb enhancer. (F) Percentage PD1-positive cells as well as PD1 MFI in CRISPRa-Jurkats targeting the *PDCD1* upstream enhancer (gated on live cells). Paired two-tailed t-test. *) *p*-value <0.05. (G) *TCF7* gene locus with pseudo bulk ATAC-seq signal from core clusters with peak-to-gene links. Highlighted is the CRISPRi/a-targeted -27kb enhancer upstream of the TSS. (H) Flow cytometry plot showing TCF1 expression in Jurkat T cells transduced with dCas9-ZIM3 and electroporated with a control gRNA or a gRNA targeting the -27 kb enhancer at the *TCF7* locus. (I) Percentage TCF1-positive cells as well as TCF1 MFI in CRISPRi-Jurkats targeting the *TCF7* upstream enhancer (gated on live cells). Paired two-tailed t-test. *) p-value <0.05; **) p-value <0.01.

## Discussion

In this manuscript we uncover an underlying core chromatin program for functional and dysfunctional human TIL states with diverse enhancer landscapes regulating cell-state-specific genes. We identified unifying chromatin signatures for cytotoxic and memory CD8^+^ T cells, and a progressive chromatin-based trajectory from precursor exhausted to terminally exhausted TILs. We support these findings by reconciling marker gene activity, surface PD1 expression, non-tumor control samples, and published gene expression signatures. Defining this core pathway to dysfunction would not have been possible without probing many patients over several entities. As an example, by solely focusing on the HNSCC samples that showed less dysfunctional TILs, we might have missed the extreme exhaustion states and heat shock signatures we derived from the remaining entities. Thus, we cover a wide breath of T cell states with our dataset, from exhausted to fully functional T cells according to flow cytometry analysis, which allows a comprehensive view on the gradients of T cell (dys)functional states in the context of cancer.

Importantly, we discover an additional level of heterogeneity among chronically activated T cells in the tumor that is likely driven by the tumor tissue of origin. Several thousand chromatin regions were significantly more accessible in T cells from either HCC, ccRCC, or BCC. We and others recently described tissue-specific epigenome remodelling of immune cells under homeostatic conditions^33, 34^, and by side-to-side comparison of several entities we now also observe cancer-type-specific chromatin remodelling in TILs. As this was more prominent with increasing dysfunction, a provocative hypothesis would be that the T cells’ chromatin landscape is more permissive for tissue effects during chronic activation, which could be highly relevant in the entity-specific targeting of T cells by immunotherapy. The converse conclusion of this heterogeneity is that analysis of TILs from a single entity contains tissue-specific chromatin patterns. This may therefore prevent identification of general genes that drive (dys-)function independent of the T cells’ residing tissue. Here, our data extends previous studies^20^ by providing a core set of genes for functional and dysfunctional T cell states captured over several entities on the single cell level.

Mapping the activity of transcription factors as upstream regulators of cell states is pivotal to understand molecular drivers of cellular function. In this regard, our study contributes new insights into the gene regulation of TILs by identifying enriched transcription factor motifs as well as transcription factor footprints at cell-state-specific open chromatin regions. Specifically, we discovered a loss of KLF and ETS:RUNX activity in T_EX_, while an intense heat shock signature was dominating both cytotoxic and dysfunctional TILs, suggesting that heat shock responses exert additive chromatin remodelling to other T cell states. In light of the sustained positive effect of heat^50^ on tumor control by T cells this observation merits further investigation of the underlying molecular mechanisms of heat shock proteins on T cell gene-regulatory cues. Further, our analysis suggests an involvement of different nuclear receptors and octamer transcription factors in shaping T cell dysfunction in the tumor. In this regard, we found increased gene activity or transcription factor motif enrichment in T_EX_ clusters for *VDR, RXR, Erra*, and *POU2AF1*, which merits further investigations.

The discovery of precursor exhausted T cells (also termed stem-like or exhausted progenitor cells), which give rise to the terminally differentiated exhausted cells that mediate pathogen/tumor eradication, spawned high interest in their molecular and cellular properties. A challenging issue in the human system remains clarifying the development and role of precursor exhausted T cells in tumors. In this study, we define a cluster that shows activity of the T_PEX_ markers *TCF7* and *CXCR5* and is enriched in the signature of T_PEX_ derived from B16 tumors and chronic LCMV infection models. Yet, the signature was discrepant to a precursor signature described recently^18^. The respective signature was derived from a mouse model of chronic infection at early timepoints of disease that markedly differs from our T_PEX_-like cluster from established human tumors, which could explain the observed discrepancy and urges probing T cell chromatin landscapes at high resolution form human chronic infections. We speculate that the core T_PEX_ (cluster 3, **Figure 4A**) represents activated but functional T cells that can progress in their chromatin signature to terminally dysfunctional cells. Several studies identified a TCF1^+^ population of T cells also in human tumors, which was associated with self-renewing capacity of TILs and good response to checkpoint blockade^5, 6^. In line with these reports, we observe an expansion of the T_EX_ cluster upon checkpoint blockade, indicating an expansion of terminally T_EX_ from precursor populations. However, whether the cluster in our analysis comprises tissue-resident or infiltrating bystander CD8^+^ T cells, infiltrating antigen-specific T cells that expand from a T_PEX_ outside of the tumor, or a mixture thereof, needs further clarification.

Besides describing modes of gene-regulation, we were able to map enhancer-promoter links in human TILs by integrating our chromatin data with scRNA-seq. We confirmed that our predicted interactions recapitulate previously described enhancer-promoter relationships and are enriched in physical chromatin contact data. We further validated a direct impact on target gene transcription of selected enhancers by targeting them with potent CRISPR-based activators and repressors. Importantly, we could prioritize cell-state-specific regulators by ATAC-seq-based super enhancer analysis. This approach was supported by top-ranking well-described genes such as *TOX* for terminally dysfunctional cells^24, 25, 26^, *TBX21* and *ZEB2* for cytotoxic T cells^51, 52^, and *TCF7* for precursor dysfunctional T cells^4, 9, 53^, and yielded many putative key genes that are unknown in their contribution to T cell states in the human system so far. Thus, we provide a unique resource comprising ranked candidate genes and connected enhancers of distinct T cell states that will advance the detailed dissection of T cell (dys-)function in the human system in the future. Importantly, our data comprehensively addresses the heterogeneity of TILs and therefore extends current knowledge on enhancer regulation in human CD8^+^ T cell subsets derived from bulk data^17^. This is illustrated by the manifold key genes and state-specific enhancers within the different TIL clusters, which was not resolved in previous studies.

Our work has several limitations. First, we analyze only four entities, and therefore do not cover impacts of all possible microenvironments, cancer subtypes, and treatments on TIL chromatin landscapes. Nevertheless, our data comprises 47 samples from 16 donors (58 samples from 23 donors including PBMC controls), and the resulting data could be successfully aligned to generate general chromatin signatures in TILs. With the vastly increasing throughput of single-cell profiling methods, future studies incorporating more samples and entities have the potential to work out even more fine-grained classification of human TIL states in relation to additional parameters such as residing tissue type, molecular cancer (sub-)classification, and treatment. Second, the provided enhancer-promoter interactions rely on computational models. Input requirements impede chromatin conformation capture-based assays on primary human TILs, but technological advances might overcome this hurdle and provide an additional experimental layer of information on gene-regulation of human TILs in the future. Nevertheless, we could successfully validate selected interactions via both up- or downregulation of predicted target genes by enhancer manipulations. Third, our ATAC-seq data does not recapitulate clonal relationships between cells as it is possible by single-cell T cell receptor analysis. Hence, we cannot follow the chromatin dynamics of single T cells by correlating discrete chromatin states with individual TCRs. Although our derived core clusters of precursor and terminally dysfunctional TILs are enriched for matching signatures of antigen-specific T cell models of infection and cancer, future studies could directly address clonal relationships by adapting multi-omics assays to capture TCR sequences along chromatin accessibility profiles.

Our study has several therapeutic implications: to begin with, we prioritize epigenetically remodeled, potential drivers and mediators of T cell (dys-)functional states that are common over several entities. Hence, overexpressing or deleting these genes represent attractive starting points to improve adoptive T cell products such as CAR T cells. Further, and in contrast to complete gene ablation using classical CRISPR knockouts, the inhibition/activation via enhancers enables tuning of gene expression, which would be desirable for certain applications. As an example, the balance of transcription factors in T cells determines the differentiation fate of lymphocytes as exemplified by T-bet and Eomes^54^, hence complete ablation of any factor might not lead to favorable T cell phenotypes. Similar, TOX was described to be not only a driver of T cell exhaustion, but also necessary for tumor and virus control, so the simple deletion of this gene did not improve T cell function in *vivo*^24, 25, 26^. Hence, tuning the expression levels of such master regulators via CRISPRi/a of linked enhancers provides a new perspective to improve immune cell function. As most genes are regulated by several enhancers that account only for a fraction of the transcriptional output as shown by our CRISPRi/a experiments, activating or repressing several non-coding elements may represent a valid strategy to fine tune expression levels. Additionally, as enhancers bind signal-specific transcription factors it may be possible to interfere with signal-specific gene-regulation by targeting the respective enhancer. Finally, we uncover many state-specific enhancers in TILs that distinguish functional from dysfunctional cells in the tumor. Such enhancers provide a unique opportunity for perturbations to abrogate expression of genes that are harmful in the tumor-context but beneficial otherwise. In the case for PD1, in mice it was shown that a certain PD1 expression is required for optimal memory formation^55^. However, strong expression in the tumor is detrimental for TILs, so interference with the exhaustion-specific enhancers therefore could reduce PD1 on TILs in the tumor context, but leave beneficial expression patterns outside of the exhaustion context intact. Our single-cell based deconvolution of TIL heterogeneity is therefore ideally suited to identify targetable regulatory elements that are context-restricted and specific to functional or dysfunctional T cell states as demonstrated by our manipulation of a T_MEM_-specific enhancer at the *TCF7* locus.

In summary, our study advances the research on human tumor-related T cell (dys-)function by defining general gene-regulatory states and a core trajectory to human T cell exhaustion. By integrated analysis of single-cell chromatin and RNA profiling we mapped putative enhancer-promoter interactions of cell-state-specific genes. Our results open new avenues for detailed studies of key non-coding gene-regulatory elements involved in steering T cell (dys-)function, and provides potential for CRISPRi/a mediated enhancer editing for in-depth studies of T cell gene regulatory cues and immunotherapeutic purposes.

## Acknowledgements

Sequencing and demultiplexing was conducted at the Core Unit of the Leibniz Institute for Immunotherapy (LIT), University Regensburg and University Medical Center Regensburg, Germany We thank the members of the LIT FACS Core Facility for their excellent technical support. We thank Manuela Kovács-Sautter and Brigitte Wild for their excellent technical assistance. We thank Charles Gersbach for providing the plasmid used for the cloning of our CRISPRi/a system. J.M.W. received funding through grant WE4675/3-1 from the Deutsche Forschungsgemeinschaft (DFG), Bonn, Germany. I.U. and P.J.S. are supported by Else Kröner Fresenius Foundation. C.S. received funding through grant SCHM 3397/3-1 from the Deutsche Forschungsgemeinschaft (DFG), Bonn, Germany.

## Author contributions

Conceptualization, C.S.; Methodology, C.S., D.R., E.R.F.; Formal Analysis, M.S., C.S., D.R.; Investigation, D.R., M.S., E.R.F.; Resources, A.R.A., K.S., R.M., F.W., M.K., B.B., I.U., J.M.W., P.J.S.; Writing – Original Draft, C.S.; Writing – Review & Editing, D.R., M.S., E.R.F, A.R.A., K.S., R.M., C.D.I., M.K., B.B., I.U., P.J.S., J.M.W., C.S.; Funding Acquisition, C.S.; Supervision, C.D.I., C.S.;

## Figure legends

**Figure S1:**
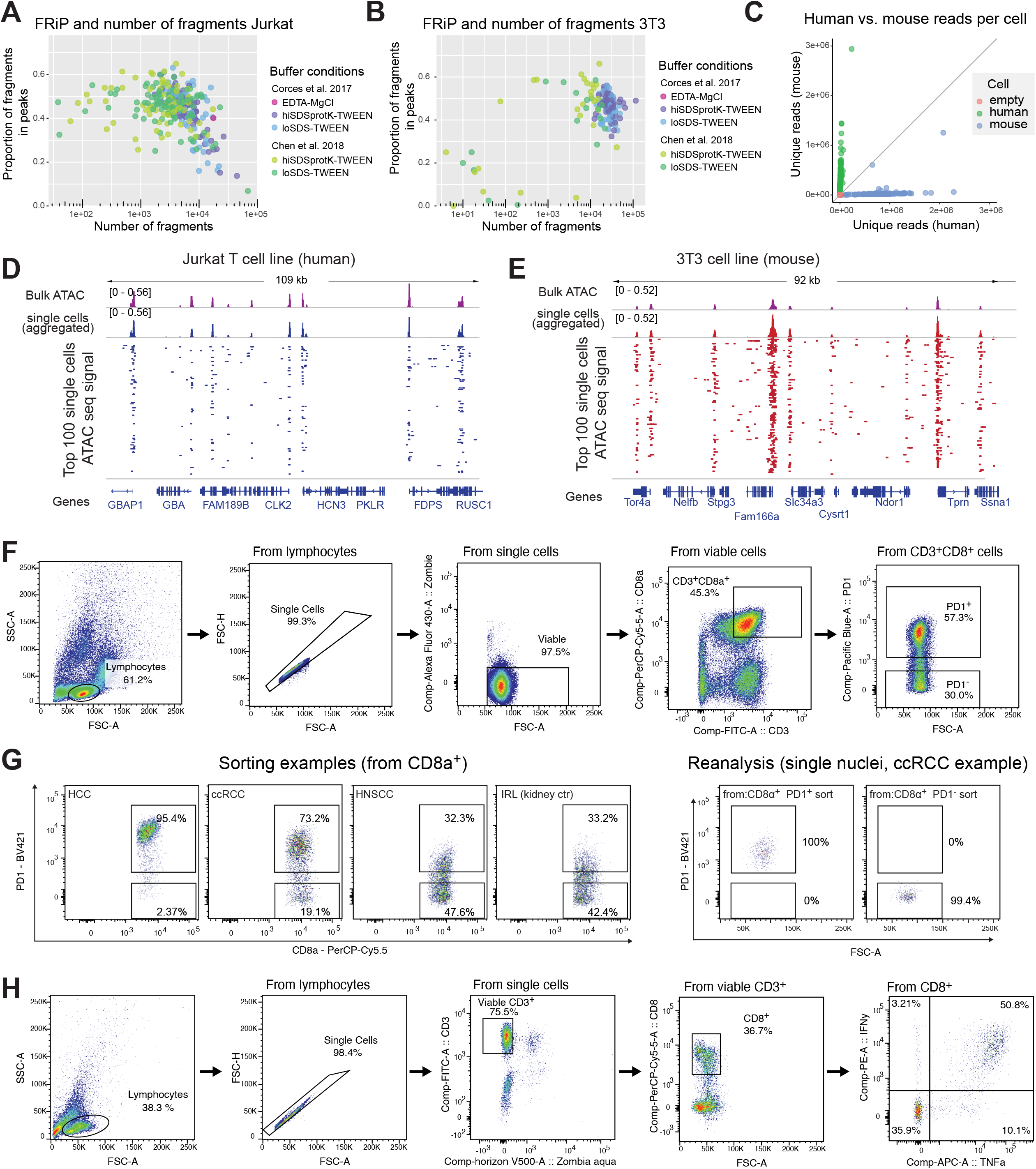
Establishment and optimization of plate-based single-cell ATAC-seq. Related to Figure 1. (A-B) Fraction of sequencing fragments in peaks and number of unique fragments with different ATAC-seq lysis buffers from Corces et al. 2017 or Chen et al. 2018 with different Tn5 neutralization conditions (“EDTA MgCl”, “high SDS and ProteinaseK”, “low SDS without ProteinaseK”) in human Jurkat (A) and murine 3T3 cells (B). (C) Evaluation of doublets arising from single-cell sorting for scATAC-seq. Unique reads aligning to the human or mouse genome from a species mixing experiment, where human Jurkat and murine 3T3 cells were mixed 1:1 and then subjected to plate-based single-cell ATAC-seq. (D-E) Genome browser tracks of bulk ATAC-seq (from 50,000 cells), merged single-cell ATAC signal, and the individual 100 single cells (top cells based on number of unique fragments) at representative genomic regions for Jurkat (D) and 3T3 cells (E). (F) Sorting strategy to sort living CD3^+^CD8^+^PD1^+^ or CD3^+^CD8^+^PD1^-^ cells for scATAC-seq of TILs and tissue controls. (G) Exemplary FACS dot plots showing PD1 expression of samples undergoing scATAC-seq (left) and sort QC (right). (H) Gating strategy for analyzing cytokine expression of CD8^+^ T cells from tumors and blood after restimulation.

**Figure S2:**
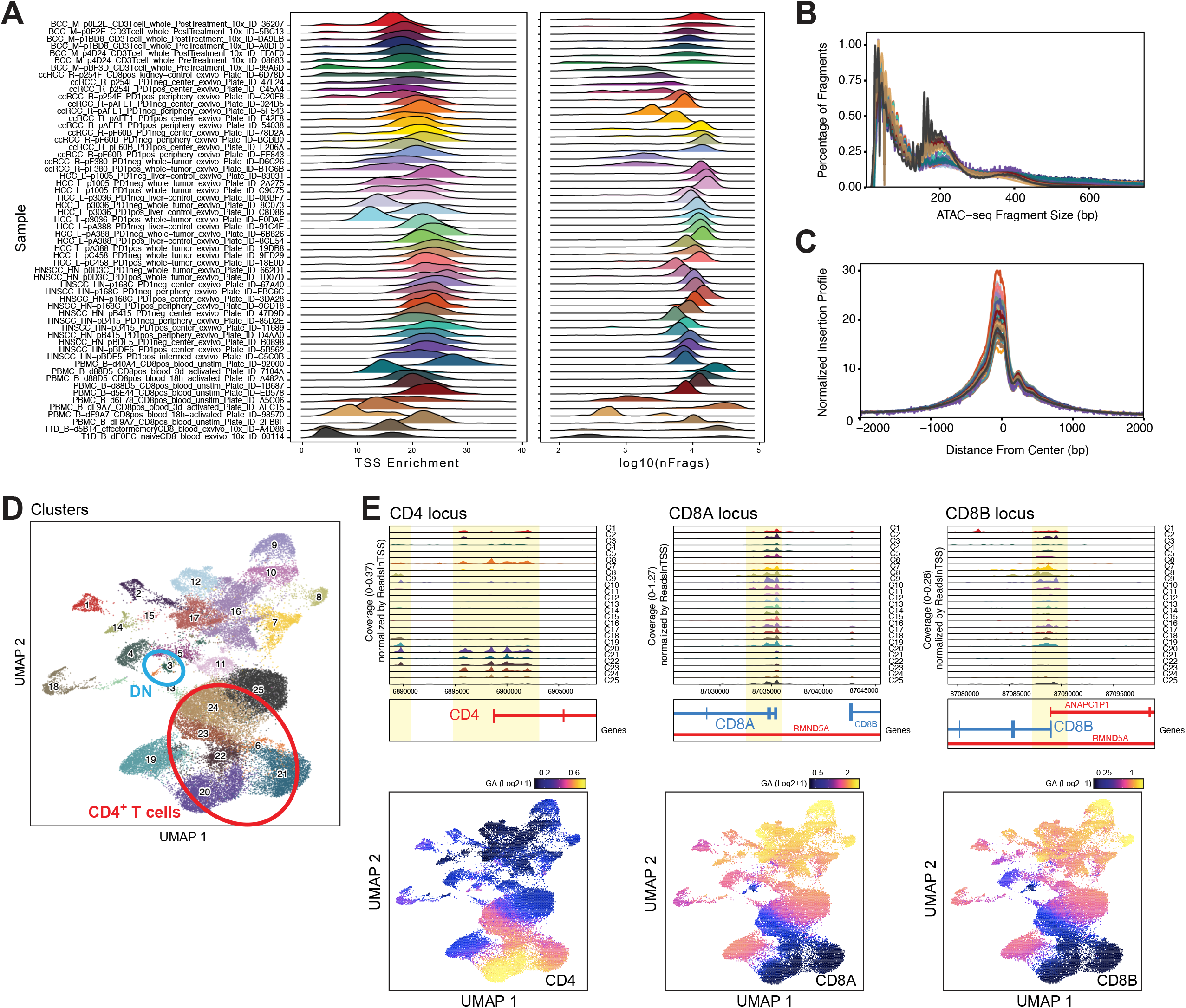
Quality control and filtering of scATAC-seq data. Related to Figure 2? (A) Transcription start site (TSS) enrichment and number of fragments (log10) for all cells in all scATAC-seq samples. (B) ATAC-seq fragment size distribution of all individual samples. (C) Sequencing fragment distribution around TSS sites (−2000 kb to +2000 kb from all annotated TSS). (D) UMAP projection and clustering of all cells in the scATAC-seq dataset. Highlighted are clusters that are CD4/CD8 double-negative (“DN”) and CD4^+^ T cells based on gene activity and pseudo-bulk ATAC-seq signals at the *CD4, CD8A*, and *CD8B* gene loci (**Figure S2E**). (E) Gene activity and pseudo-bulk ATAC-seq signals at the *CD4, CD8A*, and *CD8B* gene loci.

**Figure S3:**
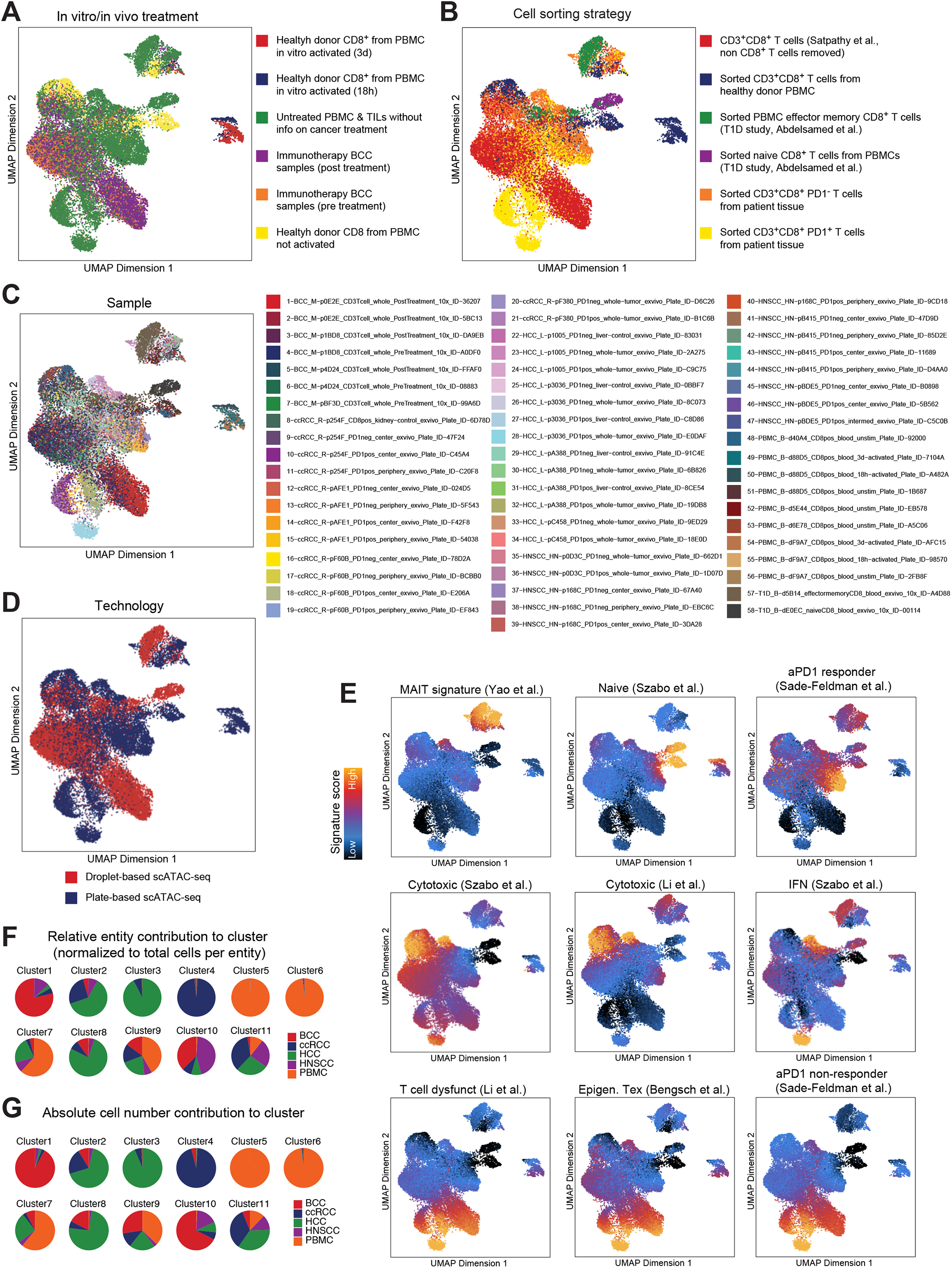
Metadata annotation of scATAC-seq data. Related to Figure 2. (A-E) UMAP projection of scATAC-seq CD8^+^ T cells. Cells are colored by *in vitro* or *in vivo* treatment (A), cell sorting/isolation prior scATAC-seq (B), individual sample identity (C), scATAC-seq technology (D), and signature enrichment score for publicly available T cell signatures (E). (F) Pie chart representation of relative contribution of entities to each scATAC-seq cluster. (G) Pie chart representation of absolute cell number contribution of entities to each scATAC-seq cluster.

**Figure S4:**
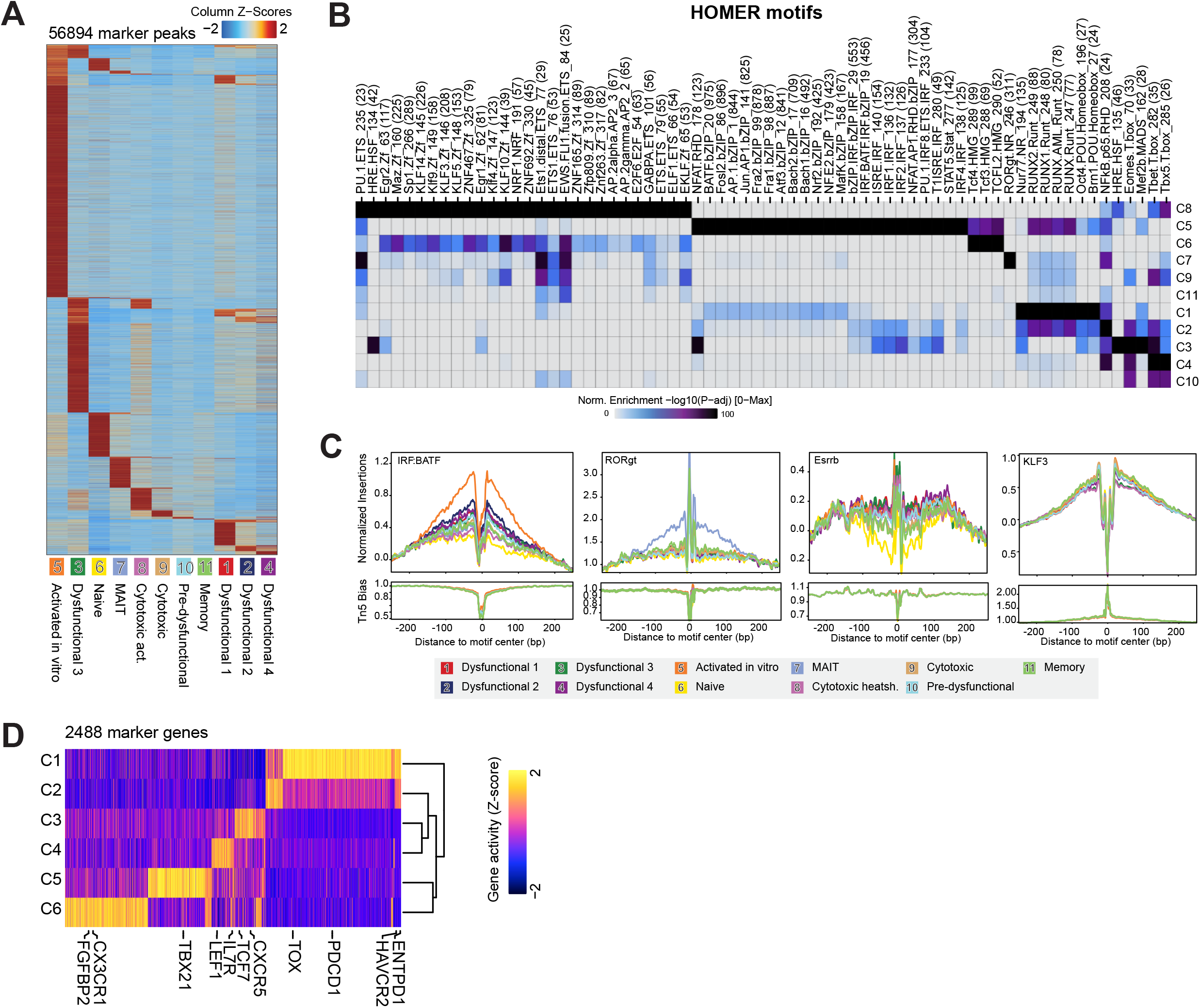
Peak-based analysis of scATAC-seq data. Related to Figure 3 and 4. (A) Heatmap showing Z-Score normalized chromatin accessibility of 56,894 marker peaks identified over all scATAC-seq clusters. (B) Enrichment of known transcription factor binding motifs (HOMER database) in cluster-specific marker peaks. (C) Transcription factor footprints of selected motifs. Shown is the Tn5 insertion preference-normalized ATAC-seq signal for each cluster at the peaks harboring the indicated TF motif. (D) Heatmap of gene activity scores of all marker genes from each cluster of the core analysis. Data is normalized (Z-Score). Statistical test for marker detection: Wilcoxon, FDR <= 0.01 & Log2FC >= 0.58.

**Figure S5:**
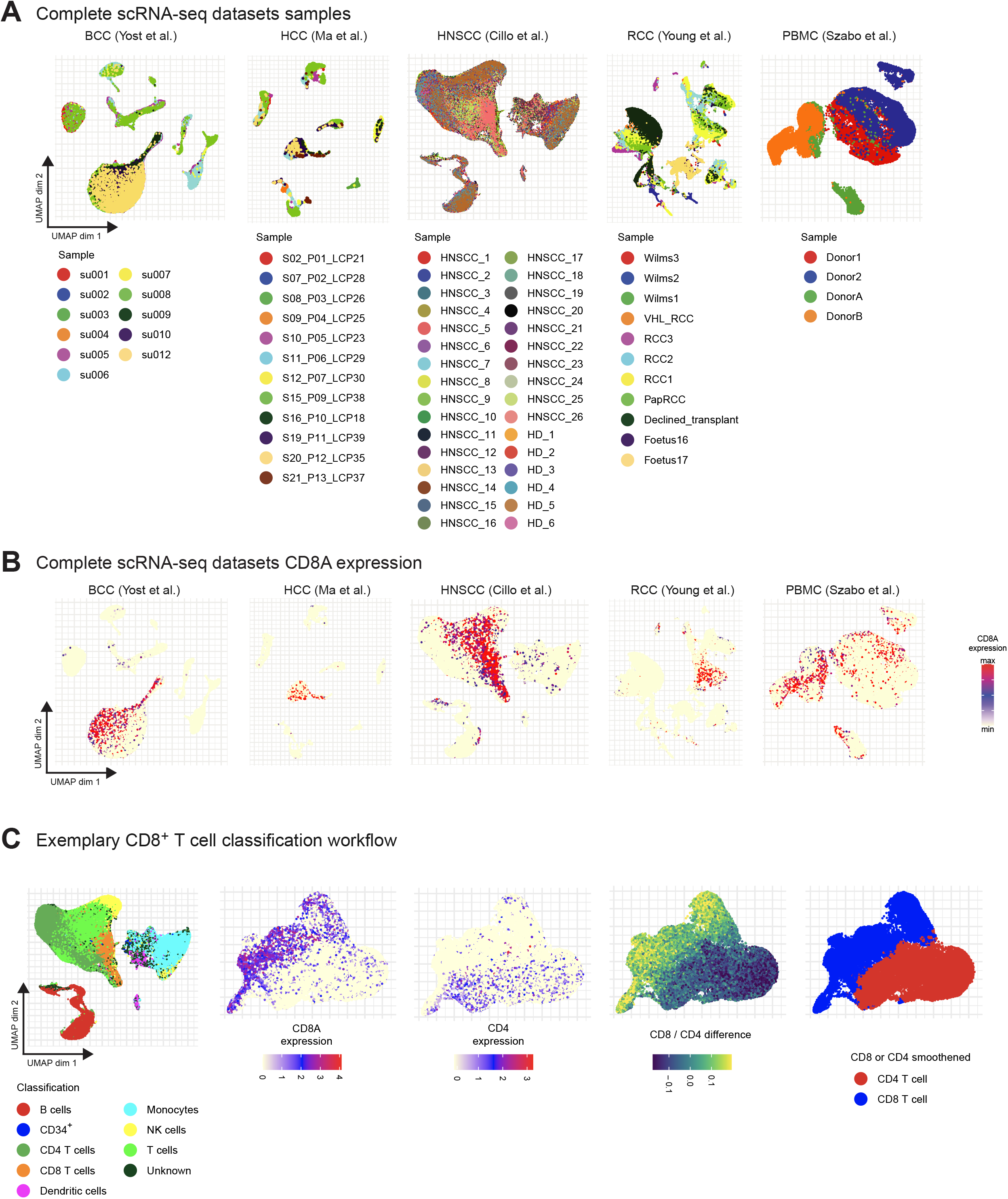
Processing and annotation of entity-matching public scRNA-seq datasets. Related to Figure 5. (A) UMAP projection and sample identity of processed single-cell RNA-seq datasets for this study. (B) CD8a expression in single cells from **Figure S5A**. (C) Exemplary workflow to remove non-CD8^+^ T cells. First, coarse cell type annotations are used to reduce data to T cells. Subsequently, a CD4 and CD8 marker gene signature score is calculated for each cell, and their difference is used to assign CD4 or CD8 cell status. Finally, cell classification is smoothed over its 20 nearest neighbors.

**Figure S6:**
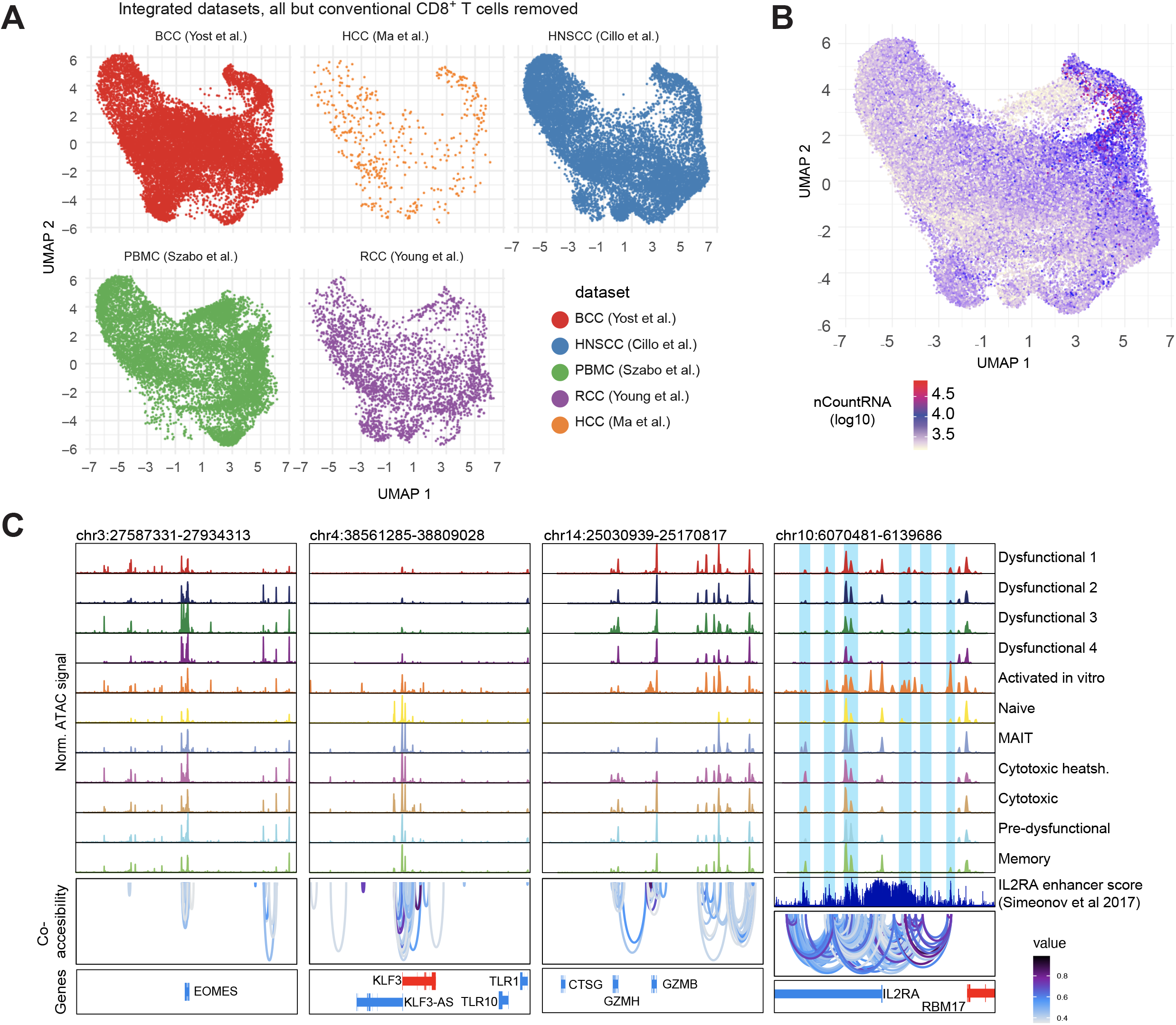
Integrated scRNA-seq. Related to Figure 5. (A) Distribution of all processed, filtered, and integrated single cells according to source of dataset. (B) Log_10_-scaled counts of unique mRNAs detected in each single cell. (C) Representative browser plot visualization of pseudo-bulk ATAC-seq signals of all clusters from **Figure 2A**, all peaks, and co-accessibility of peaks at the *EOMES, KLF3, GZMB*, and *IL2RA* loci.

## Methods

### Sample collection and cell preparation

Collection of primary tumor samples and surrounding healthy tissue from HCC, ccRCC and HNSCC tumor patients was accomplished after approval of the ethics committee (University Regensburg, reference numbers 19-1414-101, 16-355-101, 13-257-101) in accordance with the Helsinki Declaration and after signed informed consent of the patients. Peripheral blood mononuclear cells were isolated from leukocyte reduction chambers from healthy thrombocyte donors after signed informed consent (University Regensburg, reference number 13-101-0240). Primary HCC tumor samples and adjacent non-tumor liver tissue samples were collected from 4 male patients with an average age of 73.8 years (±4; range from 69 to 78) undergoing major liver resection. Primary HNSCC samples from the tumor center and the tumor periphery were collected from 3 male and 1 female patients (all HPV negative) with an average age of 57.8 years (±14; range from 43 to 71) undergoing oropharyngeal or oral cavity surgery. One patient had been treated with radiotherapy and chemotherapy previously. Primary ccRCC samples from the tumor center and the tumor periphery as well as from surrounding healthy tissue were collected after tumor resection. Immune cells from ccRCC and HNSCC tumor and tissue samples were isolated as described previously^56^. To isolate immune cells from HCC tumor and surrounding healthy liver tissue, samples were washed with HBBS medium before dissection into small fragments. After digestion with collagenase type IV and DNase I, hepatocytes were removed by filtration through a 40 µm mesh. Immune cells were then collected by ficoll gradient centrifugation. Isolated cells were frozen in RPMI containing 10 % DMSO and stored in liquid nitrogen until further use. PBMCs were isolated using Ficoll centrifugation, followed by red blood cell lysis and cryopreserved in medium containing 10% DMSO until further use.

### Cell culture

Murine NIH 3T3 cells were cultured in DMEM (GIBCO, 41965) supplemented with 10 % FCS (PAN Biotech, P30-8500) and 1 % penicillin-streptomycin. Human Jurkat cells (Clone E6-1) were cultured in RPMI (GIBCO, 11875093) containing 10 % FCS and 1 % penicillin-streptomycin (cRPMI). Cell lines were tested for mycoplasma contamination on a regular basis but not before every experiment. Lenti-X HEK293T cells (Takara Bio, 632180) were cultured in DMEM with GlutaMAX (Thermo Fisher Scientific, 31966-021) supplemented with 10% FCS (PAN Biotech, P30-8500) and 1% penicillin-streptomycin. Cells were passaged every two to three days using trypsin (Merck, T4174) to keep them at a confluency of <70%.

### Preparation of samples for FACS sorting or flow cytometry

On the day of the experiment HCC and ccRCC samples were thawed according to 10X protocol for primary/fragile cells (protocol “CG000169”, 10x genomics) followed by CD45 positive enrichment according to the manufacturer’s protocol (Miltenyi Biotec, 130-118-780). HNSCC samples were processed freshly without CD45 positive enrichment. To receive chromatin data from healthy donors as a control, PBMCs were thawed using the same procedure as for tumor samples and either sorted directly (“resting”) or activated for 18 hours (“18 hours activated”) or 3 days (“3d activated”). In short, 100,000 cells per well were activated with anti-human TransAct (1:100, Miltenyi Biotec 130-111-160) in T cell medium (10 % FCS, 1 % penicillin-streptomycin, 55 µM ß-Mercaptoethanol, 1 % L-Glutamin in RPMI) supplemented with 100 units/mL IL-2 (Peprotech, 200-02) for the respective amount of time. Cell staining for FACS sorting or flow cytometry was performed as follows: Cells were stained in 1.5 mL Eppendorf tubes for cell sort or 96-well plates for flow cytometry. Antibodies were diluted in FACS (2 % FCS, 2 mM EDTA in PBS). Prior to every staining, samples were blocked with human Kiovig solution (1:20 in PBS) for 10 minutes at 4 °C. Subsequent, cells were resuspended in 50 µl antibody mastermix per million cells and surface staining was performed for 20 minutes at 4 °C. The following anti-human antibodies were used for surface staining: CD3 (HIT3a), CD8a (SK1), CD45RA (HI100), CD38 (HIT2), 41BB (4B4-1) and CD28 (CD28.2) at a dilution of 1:100; CD197 (G043H7), PD1 (NAT105) LAG3 (11C3C65) and TIM3 (F38-2E2) at a dilution of 1:50. Zombie aqua fixable viability dye (Biolegend) was used for dead cell exclusion as recommended by the manufacturer. In order to measure cytokine production of T cells derived from tumors, tissues and blood, cells were incubated with 1 x cell stimulation cocktail containing transport inhibitors (eBiosciences) diluted in T cell medium for 5 h at 37 °C. Afterwards, intracellular staining was performed using the Foxp3/Transcription Factor Staining Buffer Set (eBiosciences) according to the manufacturer’s protocol together with the following antibodies: IFNγ (4S.B3) and TNFα (MAb11) each at a dilution of 1:50. Stained cell suspensions were passed through a 40 µm filter unit before acquiring on a BD LSRII. To validate machine functionality, BD CS&T beads were used on a regular basis. Fluorescence spillover compensation for cell sorting was performed using lymphocytes stained with CD4 (OKT4) in the respective colors and with compensation beads (Miltenyi Biotec) for flow cytometry. Sorting of cells for bulk ATAC-seq and scATAC-seq was performed using a BD FACSAriaII cell sorter. Prior to sorting, cell suspensions were filtered via 30 µm pre-separation filters (Miltenyi Biotec). Post-sort quality controls were performed whenever possible. For scATAC-seq, target cells were sorted into normal Eppendorf tubes (not low-bind tubes, as this could lead to sample loss when cell numbers are limited) containing 500 μl 0.5%BSA-PBS. For bulk ATAC-seq, target cells were sorted into DNA low-bind tubes (Eppendorf, 0030108051) containing 500 μl 10%FCS-PBS. Flow cytometry data were analyzed using BD FlowJo (version 10.7.2). The gating strategy is shown in **Figure S1**.

### Bulk ATAC-seq

Chromatin accessibility mapping was performed using the ATAC-seq method as previously described^22^, with minor adaptations. Briefly, in each experiment 50,000 cells were pelleted by centrifuging for 10 min at 4 °C at 500 × g. After centrifugation, the pellet was carefully lysed in 50 µl resuspension buffer supplemented with NP-40 (Sigma), Tween-20 and Digitonin (10 mM Tris-HCl pH 7.4, 10 mM NaCl, 3 mM MgCl2, 0.1 % NP-40, 0.1 % Tween-20, 0.01 % Digitonin) and incubated for 3 minutes on ice. Then, 1 mL of ice-cold resuspension buffer supplemented with Tween-20 was added, and the sample was centrifuged at 4 °C at 500 × g for 10 minutes. The supernatant was discarded, and the cell pellet was carefully resuspended in the transposition reaction (25 µl 2 × TD buffer (Illumina), 2.5 µl TDE1 (Illumina), 16.5 µl PBS, 5 µl nuclease-free water, 0.5 µl 1 % Digitonin (Promega), 0.5 µl 10 % Tween-20 (Sigma)) for 30 min at 37 °C on a shaker at 1000 rpm. Following DNA purification with the Clean and Concentrator-5 kit (Zymo) eluting in 23 µl, 2 µl of the eluted DNA was used in a quantitative 10 µL PCR reaction (1.25 µM forward and reverse custom Nextera primers^22^, 1 × SYBR green final concentration) to estimate the optimum number of amplification cycles with the following program 72 °C 5 min; 98 °C 30 s; 25 cycles: 98 °C 10 s, 63 °C 30 s, 72 °C 1 min; The final amplification of the library was carried out using the same PCR program and the number of cycles according to the Cq value of the qPCR. Library amplification using custom Nextera primers was followed by SPRI size selection with AmpureXP beads to exclude fragments larger than 1,200 bp. DNA concentration was measured with a Qubit fluorometer (Life Technologies). The libraries were sequenced using the Illumina NextSeq 550 platform. Equimolar amounts of each library were paired-end sequenced (2 × 38 bp) on a NextSeq 550 instrument (Illumina).

### Optimization of plate-based scATAC-seq

To establish and optimize plate-based scATAC-seq as published by Chen et al. 2018, human Jurkat T cell line and murine NIH 3T3 cell line was mixed 1:1 and 50,000 viable cells were FACS sorted as described previously by excluding 7-AAD positive cells. Subsequently, cells were pelleted by centrifuging for 5 minutes at 500 x g at 4 °C and washed twice with pre-chilled 1 × PBS. After discarding the supernatant, cells were either resuspended in 50 µl tagmentation mix “Chen et al. 2018” consisting of high salt TD buffer (33 mM Tris-HCl pH 7.8, 66 mM KCl, 10 mM MgCl2, 16 % dimethylformamide) supplemented with 0.5 µl 1 % Digitonin (Promega) and 5 µl Tn5 (Illumina) in nuclease-free water or in 50 µl tagmentation mix “Corces et al. 2017” consisting of 1 × low salt TD buffer (2 × buffer: 20 mM Tris-HCl pH 7.6, 10 mM MgCl2, 20 % dimethylformamide) supplemented with 16.5 µl PBS, 0.5 µl 1 % Digitonin and 5 µl Tn5. The tagmentation reaction was then incubated for 30 min at 37 °C on a shaker at 800 rpm. Thereafter, the reaction was stopped by adding 50 µl ice-cold tagmentation stop buffer (10 mM Tris-HCl pH 8.0, 20 mM EDTA pH 8.0 in nuclease-free water) and incubated for 10 minutes on ice. Subsequently, 100-300 µl 0.5 % BSA-PBS was added and the mix was transferred to a FACS tube where nuclei were stained with 0.2 µg/mL propidiumiodide (PI, Sigma) for 15 minutes at 4 °C in the dark. Single nuclei were sorted into prepared 384-well plates containing 2 µl 2 × reverse crosslinking buffer (RCB) and 2 µl of 10 µM forward and reverse custom Nextera index primer mix^22^. To test the performance of different reverse crosslinking buffers, nuclei which were tagmented with “Chen et al. 2018” tagmentation mix were either sorted into plates containing RCB buffer for condition “hiSDSprotK-TWEEN” (2 x RCB: 100 mM Tris-HCl pH 8, 100 mM NaCl, 40 µg/mL Proteinase K (NEB), 0.4 % SDS (Sigma) in nuclease-free water) or into plates containing RCB buffer for condition “loSDS-TWEEN” (2 x RCB: 100 mM Tris-HCl pH 8, 100 mM NaCl, 0.04 % SDS in nuclease-free water). The same was done for nuclei which were tagmented with “Corces et al. 2017”. For a fifth condition, parts of the “Corces et al. 2017” nuclei were sorted into plates containing RCB buffer for condition “EDTA-MgCl” (2 x RCB: 50 mM EDTA in nuclease-free water). After sorting, nuclei were spun down and plates were incubated for 30 min at 65 °C with lid temperature set to 105 °C to perform Tn5 release and proteinase K digestion (where applicable). To quench SDS in RCB buffers containing SDS, 4 µl of 10 % TWEEN-20 were added to every well. To wells containing RCB buffer with EDTA, 4 µl of 25 mM MgCl_2_ were added. To estimate the optimum number of amplification cycles, a quantitative PCR reaction (10 µl 2 x PCR master mix (NEB), 1 x SYBR green (Thermo Fisher), filled up to 20 µl per well) was carried out using 8 nuclei and the following program: 72 °C 5 min; 98 °C 30 s; 30 cycles: 98 °C 10 s, 63 °C 30 s, 72 °C 1 min. The final amplification was performed using the same settings and the number of cycles according to the average of the Cq values of the 8 qPCRs. After pooling all reactions, DNA was purified using the MinElute Purification Kit (Qiagen) followed by SPRI size selection (lower cut-off, 1.8 x) with AmpureXP beads to exclude fragments smaller than 150 bp. DNA concentration was measured with a Qubit fluorometer (Life Technologies) and average fragment size was calculated using the Agilent Bioanalyzer. Equimolar amounts of each library were paired-end sequenced (2*38 bp, 75 cycles) on a NextSeq 550 instrument (Illumina) using the NextSeq 500/550 High Output Kit v2.5 (Illumina).

### scATAC-seq of primary samples

Human T cells from tumors, surrounding tissue and blood were isolated, pre-enriched, stained and sorted as described before. Gates and quality control of sort are shown in **Figure S1**. To receive an approximate information about the exhaustion level of cells derived from tumors and surrounding tissue on the proteomic level, either Dead^-^CD3^+^CD8^+^PD1^+^ or Dead^-^CD3^+^CD8^+^PD1^-^ were sorted into prepared Eppendorf tubes. Examples for different PD1 expression levels of TILs are shown in **Figure S1G**. As a control, either rested or activated Dead^-^CD3^+^CD8^+^ cells were sorted from peripheral blood of healthy donors. scATAC-seq of the sorted cells was performed as described in the section above but with minor adaptions. After cell sorting, the washing step was skipped and instead steps from 10x Genomics protocol (protocol “CG000169”, 10x Genomics) for low cell input nuclei isolation were implemented: after centrifugation (300 x g, 5 min, 4 °C) cells were lysed by adding 100 µl of chilled lysis buffer (10 mM Tris-HCl pH 7.4, 10 mM NaCl, 3 mM MgCl2, 0.1 % TWEEN-20, 0.1 % NP-40, 0.01 % Digitonin, 1 % BSA in nuclease-free water) and pipette mixed followed by incubation on ice for 3 minutes. Subsequently, 300 µl pre-cooled wash buffer (10 mM Tris-HCl pH 7.4, 10 mM NaCl, 3 mM MgCl2, 0.1 % TWEEN-20, 1 % BSA in nuclease-free water) was added and pipette mixed 5 times. After centrifugation (500 x g, 5 min, 4 °C) 50 µl tagmentation mix according to Corces et al. 2017 with minor adaptions (1 × TD buffer as described previously, 16.5 µl PBS, 0.5 µl 1 % Digitonin, 0.5 µl 10 % TWEEN-20, 2.5 µl Tn5 in nuclease-free water) was added and DNA was tagmented for 30 minutes at 37 °C on a shaker set to 800 rpm. After stopping the tagmentation reaction and staining nuclei with PI, single nuclei were sorted into 384-well plates containing 2 µl of the “loSDS” 2 x RCB buffer (100 mM Tris-HCl pH 8.0, 100 mM NaCl, 0.04 % SDS) as well as forward and reverse custom Nextera index primer mix. Tn5 release was performed via incubation at 55°C for 30 minutes, followed by quenching SDS with 4 µl of 4 % TWEEN-20. Final amplification PCR was performed as described before by adding 8 µl of 2 x PCR MM (NEB) with number of cycles according to qPCR (usually Cq values were around 20). Sequencing of libraries was performed as described previously.

### Pre-processing and analysis of bulk ATAC-seq data

Demultiplexing was performed with bcl2fastq using the parameters: --barcode-mismatches 1. ATAC-seq reads were trimmed using Skewer^57^ and aligned to the hg19 assembly of the human genome using Bowtie2^58^ with the ‘--very-sensitive’ parameter and a maximum fragment length of 2,000 bp. Duplicate and unpaired reads were removed using the sambamba^59^ ‘markdup’ command, and reads with mapping quality >30 and alignment to the nuclear genome were kept. All downstream analyses were performed on these filtered reads. All reads aligning to the “+” strand were offset by +4 bp, and all reads aligning to the “–” strand were offset −5 bp to represent the center of the transposon binding event as described previously^60^. For visualization purposes only, coverage files from filtered bam files were produced using bedtools^61^ genomeCoverageBed, each position was normalized by dividing to the total library size and multiplying by 10^6^, followed by conversion to bigWig format using the bedGraphToBigWig command from the UCSC genome browser tools.

### Pre-processing of single-cell ATAC-seq data

Demultiplexing was performed with bcl2fastq using the parameters: --barcode-mismatches 0. Demultiplexed single-cell sequencing reads belonging to the same sample were combined and each read was brought into a format starting with “@” + “barcode” + “:” + “read_name” so it could be processed with the SnapATAC pipeline^62^ (mapping to the hg19 genome using bowtie2 with the option “—maxins 2000”). Duplicates of mapped bamfiles were marked and removed using sambamba to keep only unique and paired reads with a quality ≥ 30.

### Analysis of single-cell ATAC-seq data

Pre-processed ATAC-seq data from droplet-based scATAC-seq processed with cellranger or filtered .bam files from plate-based scATAC-seq were read with the ArchR R package^37^ retaining cell barcodes with at least 100 fragments per cell and a TSS enrichment score ≥ 3. Doublets were identified and filtered (‘addDoubletScores’ and ‘filterDoublets’). After quality evaluation, further filtering was applied, keeping only cell barcodes with at least 1000 fragments per cell and a TSS enrichment score ≥ 10. To filter non CD8^+^ T cells dimensionality reduction was performed with the ‘addIterativeLSI’ method (iterations = 4, res = 2, maxClusters = 6, varFeatures = 25000, dimsToUse = c(1:24), LSI method: “log(tf-idf)”, scaleDims=TRUE). Inspection of explained variances suggested to further use dimensions 1-21 for downstream analyses. Iterative LSI was then followed by clustering (‘addClusters’, res = 1, dimsToUse =c(1:21), scaleDims=TRUE) and UMAP dimensionality reduction (nNeighbors = 30, metric = cosine, minDist = 0.3). Based on gene-activity scores of *CD8A, CD8B*, and *CD4* loci, non-CD8^+^ clusters were removed from the dataset. The remaining cells were again subjected to iterative LSI dimensionality reduction (iterations = 3, res = c(0.1, 0,4), varFeatures = 25000, dimsToUse= c(1:24), LSI method: “log(tf-idf)”, scaleDims=TRUE). Inspection of explained variances suggested to further use dimensions 1-11 for downstream analyses. Clustering (addClusters, res = 1, dimsToUse =c(1:11), scaleDims=TRUE) was performed, and batch effects arising from the scATAC technology (plate vs droplet-based processing) were removed using Harmony^63^. UMAP dimensionality reduction was performed to visualize data. To estimate gene activity the ArchR gene activity scores was used and activity scores for visualization were imputed using MAGIC^64^. Marker genes were identified using the ‘getMarkerFeatures’ and ‘getMarkers’ functions with a foldchange of 1.5 (wilcoxon test, cutOff “FDR <= 0.01 & Log2FC >= 0.58”). Single-cell chromatin accessibility data were used to generate pseudobulk replicates from scATAC-seq datasets (ArchR functions ‘addGroupCoverages’ and ‘addReproduciblePeakSet’) for peak calling with macs2^65^. Marker peaks for each cluster for peak-based analysis were called using the ‘getMarkerFeatures’ function on the ArchR peakmatrix (wilcoxon test, cutOff = “FDR <= 0.01 & Log2FC >= 1”). A background peak set controlling for total accessibility and GC-content was generated using ‘addBgdPeaks’. ChromVAR^31^ was run with ‘addDeviationsMatrix’, using the HOMER motif set^66^ to calculate enrichment of chromatin accessibility at different TF motif sequences in single cells. To visualize motif deviations, scores were imputed using MAGIC. Transcription factor footprintig was performed and visualized using the ArchR functions ‘getFootprints’ and ‘plotFootprints’ (flank = 250, flankNorm = 50, normMethod = “subtract”, smoothWindow = 6”). With Applying the same cutoffs (“FDR <= 0.01 & Log2FC >= 1”), pair-wise comparisons for marker peak detection were performed, and the resulting peak sets were analyzed for *de novo* motif enrichment using HOMER’s ‘findMotifsGenome.pl’ function. Co-accessibility was computed with the ArchR function ‘addCoAccessibility’.

### Analyis of core ATAC-seq profiles

The scATAC-seq dataset was subsetted by removing all PBMC samples and the MAIT cell cluster. Dimensionality reduction was performed with the ‘addIterativeLSI’ method (iterations = 3, res = c(0.1, 0.4), varFeatures = 25000, dimsToUse = c(1:24), LSI method: “log(tf-idf)”, scaleDims=FALSE), using scaleDims=FALSE with the purpose to reduce stronger sample-specific biases as we aimed to generate uniform ATAC-seq signature for archetypical T cell chromatin landscapes. The first dimension was filtered due to high correlation with sequencing depth (> 0.75) according to ArchR’s standard settings. Inspection of explained variances suggested to further use the following 13 dimensions for downstream analyses. Next, harmonization for technology (10x droplet based scATAC-seq vs. plate-based approach) and entity, clustering (‘addClusters’, res = 0.3), and UMAP dimensionality reduction were performed. Marker genes, peak calling, marker peak identification, and ChromVAR motif deviations were performed as in the previous section “Analysis of single-cell ATAC-seq data”.

### Enrichment of signatures in ATAC-seq clusters

Published signatures (**Table S7**, containing also preprocessed signatures ^8^) were used to derive the expression of genes in these lists in single cells (“signatures”) by using the ‘addModuleScore’ function from ArchR. Next, the median module score for each signature in each cluster was collected and plotted using the ggheatmap package, normalizing by the “percentize” function.

### Pseudotime trajectory analysis

To analyze dynamics of peak accessibility, transcription factor activity, and gene activity along an hypothetical time axis identified by progressive chromatin changes from a precursor exhausted to a terminally dysfunctional T cell population, pseudotime analysis was performed. The harmony-corrected ArchR object of the core analysis was subjected to “addTrajectory” using the following user-defined trajectory as a guide: “Cluster 3” → “Cluster 2” → “Cluster 1”. peak accessibility, ChromVAR deviation scores, and gene activity scores were correlated throughout pseudotime using ‘getTrajectory’ and ‘plotTrajectoryHeatmap’ using standard parameters for gene activities and peak accessibility, and “varCutOff = 0.8” for the motif matrix.

### Public scRNA datasets

The following publicly available scRNA datasets were downloaded:

- HNSCC (Cillo et al. 2020, PMID: 31924475, GSE139324)
- PBMC (Szabo et al., PMID: 31624246, GSE126030)
- HCC (Ma et al. 2019, PMID: 31588021, GSE125449)
- BCC (Yost et al., PMID: 31359002, GSE123813)
- SCC (Yost et al., PMID: 31359002, GSE123813)
- RCC (Young et al., PMID: 30093597, Count data provided as Data S1 accompanying the publication)

For each dataset, cells were filtered for >200 detected genes, >1000 reads and a mitochondrial read fraction <10 %.

### Identification of CD8 T cells in scRNA datasets

For RCC, BCC, SCC and PBMC, the authors’ cell type annotations were used to select the CD8 T cell subset. For RCC, cells were additionally subsetted to the ‘Tumor_Immune’ compartment and cells from tumors classified as ‘PapRCC’ or ‘Wilms’ were excluded. For HNSCC, Garnett v 0.1.19 was used to annotate the cells with the pre-trained ‘hsPBMC’ classifier available at https://cole-trapnell-lab.github.io/garnett/classifiers/. Subsequently, all cells from the tumor samples annotated as ‘T cells’ were maintained. For HCC, cells annotated as ‘T cell’ by the authors were maintained. In a second step, a finer annotation into CD4 and CD8 T cell subsets for the T cell compartment of HNSCC and HCC was achieved based on cell type signature scores. In short, signature genes were searched based on the NSCLC PBMC signature matrix from Newman et. al. (Table S2 2a, PMID: 31061481). Genes with more than double expression value within CD8 T cells compared to CD4 T cells were selected as CD8 T cell markers and vice versa. The function ‘AddModuleScore’ from Seurat was used to obtain a CD4 and CD8 signature score for each cell using these selected markers. Cells with a CD8 score greater than the CD4 score were annotated as CD8 T cell and CD4 T cell otherwise. Subsequently, the annotation was smoothed by assigning the most common label among the 20 nearest neighbours in UMAP space to each cell and CD4 T cells were discarded.

### scRNA CD8 T cell data integration and analysis

Lower quality T cells were filtered for >1500 reads and a ribosomal read fraction <45 % and matched to scATAC-seq entities by removing squamous cell carcinoma (SCC) subset from the Yost et al. data. Each dataset was normalized (‘NormalizeData’) and variable features were identified (‘FindVariableFeatures’). Then, integration anchors between the datasets were calculated using ‘FindIntegrationAnchors(anchor.features=1000)’ and the anchorset was passed to the function ‘IntegrateData(dims=1:20)’ for dataset integration. PCA was performed followed by ‘FindNeighbors’ and ‘FindClusters’ (res=0.4) to cluster cells, followed by UMAP projection using dimensions 1:12. Cluster 8 was removed due to low count of transcripts. Data was scaled (‘ScaleData’, mode.use= ‘linear’) and marker genes were identified (‘FindAllMarkers’, min.pct =0.2, logfc.threshold=0.3).

### Integrated scATAC and scRNA-seq analysis

Single-cell ATAC and RNA sequencing data was integrated using the ArchR ‘addGeneIntegrationMatrix’ function, using the integrated scRNA-seq object and either the scATAC-seq object containing all clusters or the ‘core’ clusters, respectively. “Peak-to-gene links” (PGLs) were calculated using correlations between peak accessibility and integrated scRNA-seq expression data using ‘addPeak2GeneLinks’ function (maxDist = 400000, corCutOff = 0.75, k = 150, overlapCutOff = 0.8, predictinoCutOff = 0.35, knnIterations = 500).

### Super enhancer analysis

PGLs were calculated for the core ArchR object, and significant high-confidence PGLs (corCutOff = 0.35, FDR <= 1e-04, VarQATAC >= 0.25, VarQRNA >= 0.25) that were overlapping marker peaks of the core clusters were isolated. Next, the number of marker peaks linked to each gene via PGLs was calculated and scaled to 1 (maximum number of marker peaks). Number of linked peaks per gene were ranked and the “elbow” was determined using the ‘findElbow’ function from the R package healthcareai.

### Comparison of p2g-links with Hi-C contacts

The observed p2g-links were compared against publicly available promoter capture Hi-C data (^42^ Data S1, PCHiC_peak_matrix_cutoff5.tsv) to validate the quality of interactions. First, peak and promoter regions in the p2g-links were extended to both sides by 1 kb. Then, Hi-C contacts were filtered to a maximum distance of 300 kb between bait and promoter interacting regions to match the interaction distance constraint from p2g-links detection. Subsequently, cell type-specific Hi-C contacts were enriched by filtering for a CHICAGO score > 5 for each cell type, respectively. To obtain a similar number of specific contacts for each cell type, these Hi-C contacts were further subsampled to the minimum number of cell type-specific contacts obtained (73,921 Hi-C contacts). Eventually, the fraction of p2g-links overlapping with each of the cell type-specific Hi-C datasets was measured by searching for p2g-links that overlap matching bait and promoter interacting regions by at least 1 bp. To compute a background match fraction, random distance-matched p2g-links were generated by associating each peak center from the complete peak set (197,137 peaks) to its closest transcription start site and extending regions to both sides by 1 kb. The p2g-links in the observed dataset were binned into 100 equally spaced bins based on their widths and the frequency of each bin was calculated. Similarly, the random p2g-links were sorted into these bins and subsampled to the minimum obtained bin count. Then, p2g-links were sampled from this random p2g-link dataset according to the bin probabilities in the observed p2g-link dataset and their fraction overlapping with the cell type-specific Hi-C contacts was quantified as described above. This procedure was repeated 100 times for each cell type.

### Plasmids

For CRISPRa, the p300 core domain from plasmid pLV-dCas9-p300-P2A-PuroR^67^ was switched to a VPRmini^49^-FLAG with Gibson Assembly after BAMHI-HF (NEB, R3136S) digestion to generate pLV-dCas9-VPRmini. For CRISPRi, the p300 core domain was switched to a 2xFLAG, followed by a second digestion with AgeI-HF (NEB, cat#3552S) and AfeI (NEB, cat#R0652S) in order to clone a ZIM3 domain^48^ upstream of dCas9 and generate pLV-ZIM3-dCas9. pLV-dCas9-p300-P2A-PuroR was a gift from Charles Gersbach (Addgene plasmid #83889). All gene fragments used were ordered without adapter sequences from Twist Bioscience. The corresponding sequences can be found in the supplementary material section.

### Lentivirus Production

HEK293T cells were seeded in Opti-MEM™ Reduced Serum Medium (OPTI-MEM) with Gluta-MAX (Thermo Fisher Scientific, cat#51985-034) supplemented with 5% FCS, 1mM Sodium Pyruvate (Thermo Fisher Scientific, cat#11360-070) (cOPTI-MEM) at 1.2 million cells per well in 6-well plates and incubated overnight to reach a confluency of 90% at the moment of transfection. The transfection was performed using Lipofectamine 3000 transfection reagent (Thermo Fisher Scientific, cat#100022052) with a 2^nd^ generation lentiviral packaging system. For each well, 7 µL of lipofectamine 3000 reagent was added to 250 µL of room temperature (RT) non-supplemented OPTI-MEM medium. 1.8 µg of the corresponding CRISPRi/a transfer plasmid, 1.8 µg of psPAX2 (Addgene, #12260), 0.8 µg of pMD2.G (Addgene, 12259) and 6 µl of p3000 reagent was added to 250µl RT non-supplemented OPTIM-MEM. Mixes containing the lipofectamine and the plasmids were mixed by gentle inverted and incubated at RT for 20 min before adding it on top of the HEK293T cells. 6 hours after transfection, the medium was replaced by fresh cOPTI-MEM medium supplemented with 1x ViralBoost (Alstem, cat#VB100). Lentiviral supernatant was harvested 24h post-transfection and replaced with cOPTI-MEM supplemented with 1x ViralBoost. A second harvest was done 48h post-transfection. Both supernatants were pooled and centrifuged at 1000 x g for 5 min to remove cell debris before concentrating overnight by adding Viral Concentrator (Alstem, VC100) following the manufacturer’s instructions. The virus pellet was resuspended with ice-cold PBS to get a 100-fold concentration, and immediately used for transduction.

### CRISPR interference and activation experiments

Jurkat T cells were infected with 2.5% v/v concentrated pLV-ZIM3-dCas9 lentivirus or 5% v/v pL-dCas9-VPRmini. The next day, fresh cRPMI medium with puromycin (final concentration 2 µg/ml) was added to keep cells at a density of 0.3 million cells/ml. Three days after infection, transduction efficiency was validated by flow cytometry. For CRISPRi/a target validations, sgRNAs were introduced into the cells transiently with the 4D-Nucleofector X Unit (Lonza, cat#AAF-1003X). For each electroporation, one million cells were resuspended in 24 µl of SE buffer and mixed with 300 pmols/sgRNA before transferring to a 16-well Nucleocuvette™ well. After electroporation was performed with program CK-116, cells were kept at RT for 15 min and then transferred to a round-bottom 96-well plate in RPMI + 10 % FCS without anti-biotics to reach a final cell density of 0.5 million cells/mL. Target expression was checked by flow cytometry 24 h or 48 h after CRISPRa and CRISPRi electroporation, respectively. For *HAVRC2* enhancer-targeting, cells were pre-activated with Dynabeads human T-Activator CD3/CD28 (Thermo Fisher Scientific, cat#111.31D) for 24 h and beads were removed before electroporation. gRNA sequences are listed in **Table S8**.

## Ethics statement

Collection of primary tumor samples and surrounding healthy tissue from HCC, ccRCC and HNSCC tumor patients was accomplished after approval of the ethics committee (University Regensburg, reference numbers 19-1414-101, 16-355-101, 13-257-101) in accordance with the Helsinki Declaration and after signed informed consent of the patients. Peripheral blood mononuclear cells were isolated from leukocyte reduction chambers from healthy thrombocyte donors after signed informed consent (University Regensburg, reference number 13-101-0240).

## Data Availability

All data on primary human T cells from this study is available at the European Nucleotide Archive under the number XXX.

